# Environmental Amino Acid Sensing Regulates the Rate of ASC Translation and NLRP3 Inflammasome Assembly

**DOI:** 10.64898/2026.01.16.699988

**Authors:** Mikel D. Haggadone, Brian P. Goldspiel, Aoife O’Farrell, Nora T. Kiledjian, Montana Knight, Talia R. Smith, Elena Anderson, Víctor R. Vázquez Marrero, Mark A. Boyer, Peining Jimmy Xu, Michael Scaglione, Zachary M. Powers, Clemence Queriault, Aaron Wu, Qihua Yang, Mary X. O’Riordan, Arjun Raj, Clementina Mesaros, Crystal S. Conn, Sunny Shin, Will Bailis

## Abstract

The NOD-, LRR-, and pyrin domain-containing protein 3 (NLRP3) inflammasome is a multiprotein signaling complex that triggers pyroptotic cell death and interleukin (IL)-1 family cytokine release during infection and cell injury. Its assembly is driven by the adaptor protein, apoptosis-associated speck-like protein containing a CARD (ASC), whose filamentation forms a supramolecular speck upon NLRP3 activation to amplify inflammasome signaling. While the NLRP3 inflammasome is well appreciated as a sensor of environmental danger and damage, little is known about how homeostatic environmental factors like dietary metabolites regulate its activity. Here, we find that environmental availability of the branched-chain amino acids (BCAAs), leucine, isoleucine, and valine, controls NLRP3 inflammasome assembly. While ASC is typically viewed as a constitutively expressed, unregulated inflammasome component, we find that Toll-like receptor 4 (TLR4) activation triggers localization of ASC mRNA to the perinuclear space. Moreover, our data demonstrate that ASC undergoes TLR4-driven translational bursting from polyribosomes during inflammasome priming. This translational engagement is dependent on BCAA availability and mechanistic target of rapamycin (mTOR) activity, which regulate the kinetics of inflammasome assembly. In contrast, the translation of NLRP3 and caspase-1 is largely insensitive to these inputs. Furthermore, we find that BCAAs regulate NLRP3 inflammasome activation in both mouse and human macrophages, in the context of bacterial infection, and during lipopolysaccharide (LPS)-induced sepsis *in vivo*. Altogether, this work unveils a novel inflammasome priming event governed by the amino acid environment. These findings further highlight how the activity of proteins maintained in equilibrium like ASC can be dynamically regulated through rapid changes in mRNA translation.

## INTRODUCTION

Inflammasomes are multimeric protein complexes that initiate an inflammatory form of programmed cell death known as pyroptosis (*1–5*). By promoting the maturation of interleukin (IL)-1 family cytokines and release of alarmins during cell lysis, inflammasomes serve as critical effectors of innate immune-mediated inflammation (*6, 7*). Depending on the intensity and duration of their signaling, inflammasomes can support host defense by promoting pathogen clearance (*8–14*) or contribute to pathological inflammation during both infectious and sterile insults (*15–23*).

Among inflammasomes, the NOD-, LRR-, and pyrin domain-containing protein 3 (NLRP3) inflammasome is the most extensively studied, owing to its central role as a cytosolic sensor of cellular dysfunction and infection. Its assembly into a supramolecular signaling complex canonically requires two distinct signals. The first is a priming event triggered by pattern recognition receptor (PRR) signaling, which leads to increased NLRP3 and IL-1 expression within the cell (*24*). Once sufficiently expressed, a second pathogen- or damage-associated molecular pattern (PAMP or DAMP, respectively) activates NLRP3 (*25–30*), prompting formation of the inflammasome complex. NLRP3 recruits the adaptor protein ASC (apoptosis-associated speck-like protein containing a CARD), which rapidly oligomerizes into a large speck (*31–34*) and serves as a platform for the recruitment and activation of caspase-1 (*35, 36*), the key executioner protease responsible for downstream inflammasome effector functions (*37*). Among its substrates, including IL-1 family precursor proteins (*38, 39*), caspase-1 cleaves gasdermin D (GSDMD) and releases its N-terminal fragment, which oligomerizes and forms membrane pores that promote IL-1 cytokine release and trigger terminal cell lysis (*40*). Historically, ASC and caspase-1 have been viewed as constitutively expressed components of the inflammasome, largely insulated from biosynthetic regulation. Thus, priming has been classically understood to function by inducing NLRP3 expression and initiating post-translational modifications that license its activation (*41, 42*). Whether other priming-dependent mechanisms regulate the kinetics of inflammasome assembly, such as the localization and/or rate of inflammasome protein synthesis, remains an unresolved question.

While the NLRP3 inflammasome is well appreciated as a sensor of environmental danger and damage, much less is known about how changes in the levels of environmental homeostatic factors like metabolites impact its activity. Recent work in this area has primarily focused on how PAMP- or DAMP-induced intracellular metabolic changes tune the terminal stages of inflammasome signaling—particularly the regulation of GSDMD oligomerization and membrane insertion via reactive oxygen species signaling (*43, 44*) and lipid metabolism (*45, 46*), respectively. While these studies highlight the importance of metabolic substrates in the execution of pyroptosis, whether environmental nutrient availability is sensed and integrated as an input into inflammasome signaling, assembly, or function remains poorly understood.

Here, we identify the amino acid environment as a critical biosynthetic checkpoint that regulates NLRP3 inflammasome function. In screening the requirement of each individual amino acid for NLRP3 inflammasome activation, we found that amino acids exert distinct effects on pyroptosis, rather than macrophages responding uniformly to amino acid stress. In particular, we found that the three essential branched-chain amino acids (BCAAs)—leucine, isoleucine, and valine—act as key metabolic signals controlling the rate of NLRP3 inflammasome assembly. We then examined how BCAAs and the central environmental amino acid sensor, mechanistic target of rapamycin complex 1 (mTORC1), shape the macrophage transcriptional and translational landscape. This analysis revealed that BCAA sensing and mTORC1 signaling control rapid ASC mRNA association with polyribosomes following Toll-like receptor 4 (TLR4) activation. Using RNA-fluorescence in situ hybridization (RNA-FISH), we further demonstrate that this burst in ASC translation is spatially linked to perinuclear mRNA localization during inflammasome priming, establishing an organized translation program that is preserved during ASC speck formation. Finally, we demonstrate that the timing of mRNA translation, rather than the mere expression of inflammasome proteins, serves as a critical determinant of NLRP3 inflammasome assembly. Our data indicate that NLRP3 inflammasome priming involves not only the increased transcription of inflammasome components, but also increased translation of the adaptor protein ASC, which was previously thought to be constitutively expressed by macrophages and thus insensitive to priming events. Together, these findings uncover a new axis for homeostatic factors like metabolites governing innate immunity, positioning the amino acid environment as a key regulator of NLRP3 inflammasome activation through orchestration of a targeted translation program that seeds inflammasome complex formation.

## RESULTS

### Environmental Amino Acids Selectively and Differentially Tune Macrophage Inflammatory Cytokine Release

Innate immune sensing of PAMPs through PRRs such as the TLRs induces broad changes in gene expression and protein synthesis (*47, 48*). Accordingly, we wanted to investigate whether these energy-intensive processes engage the external macrophage metabolic environment, with a focus on amino acids as metabolites that are consumed for both energy production and protein synthesis. We looked toward the NLRP3 inflammasome as a critical driver of innate immune-mediated inflammation, and we investigated how environmental amino acid levels regulate NLRP3-triggered IL-1β production and cell death in bone marrow-derived macrophages (BMDMs). To experimentally titer global amino acid availability, we diluted a complete (i.e., control) RPMI 1640 medium with amino acid-free RPMI 1640, creating a range of amino acid levels in culture. Then, to distinguish between the acute effects of amino acid stress versus macrophage adaptation to sustained amino acid withdrawal, we implemented two experimental conditions: (1) “Immediate Restriction,” in which amino acids were limited just prior to lipopolysaccharide (LPS) stimulation, and (2) “Prolonged Restriction,” where BMDMs were cultured in amino acid-titrated medium for 20-24 hours prior to LPS treatment.

Under these conditions, BMDM viability was preserved during prolonged culture in medium containing 10% of standard amino acid levels (fig. S1A). Despite preserved viability, both the immediate and prolonged restriction of amino acids significantly impaired IL-1β release following LPS priming and subsequent NLRP3 activation with the bacterial toxin nigericin (*4*). However, only prolonged amino acid withdrawal significantly reduced NLRP3-dependent cell death in BMDMs (fig. S1, B and C). These data highlight a critical role for environmental amino acid levels in regulating NLRP3 inflammasome-driven pyroptosis.

We next investigated whether macrophages respond to prolonged amino acid restriction in a stereotyped manner or if individual amino acids exert specific effects on inflammatory cytokine release. We generated custom media in which each of the 20 protein-coding amino acids were individually restricted, with a fully supplemented medium serving as a control (Table S1). We then cultured BMDMs in each of these media for 24 hours before priming and activating the NLRP3 inflammasome with LPS and nigericin, respectively (summarized in Fig. 1A). Of note, only threonine restriction substantially compromised BMDM viability during the 20-24 hours of culture (fig. S1D). This condition was therefore excluded from downstream analyses.

**Fig. 1.**
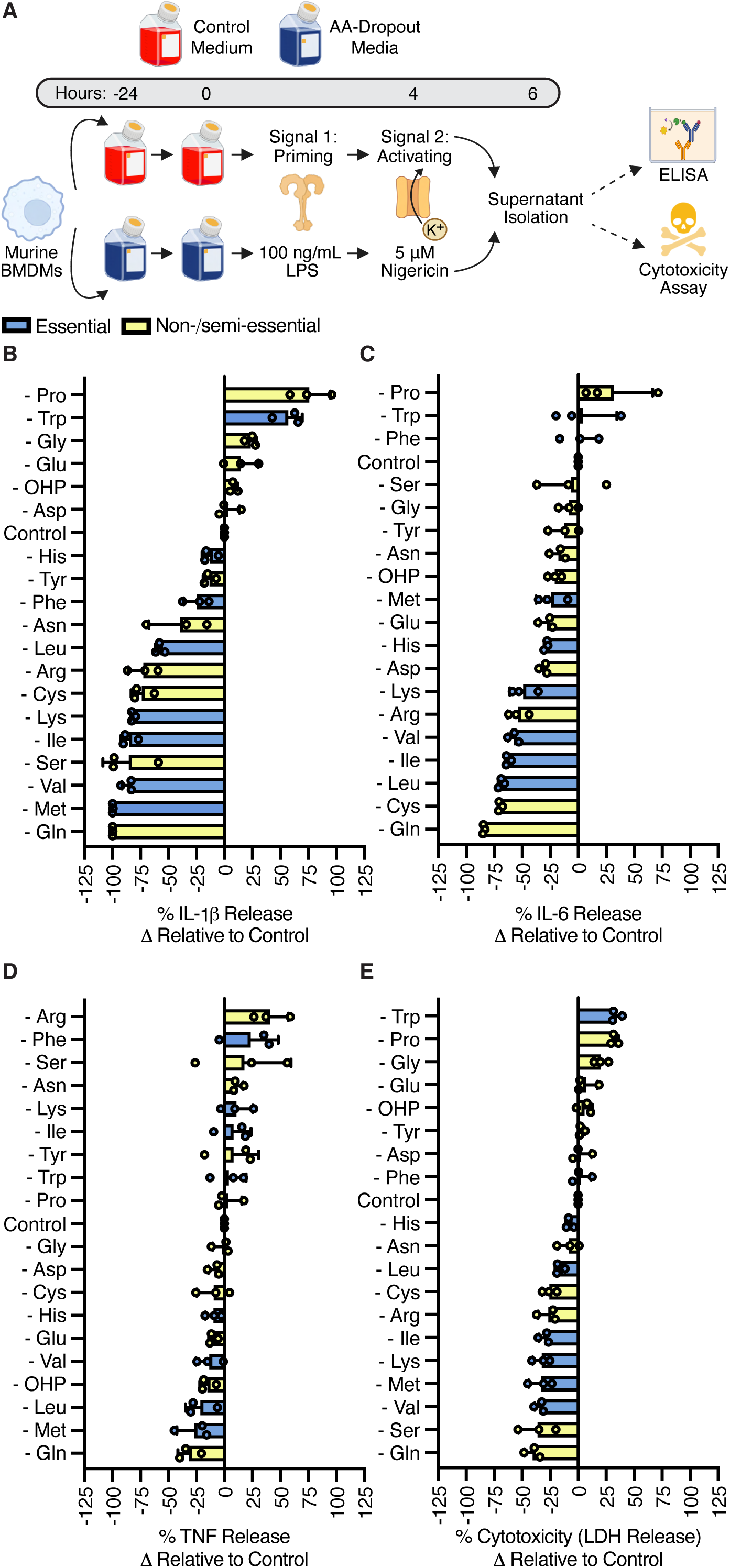
The amino acid environment tunes NLRP3 inflammasome activation and inflammatory cytokine production. (**A**) Schematic of experimental approach to limit environmental amino acids in BMDM cell culture. (**B**) to (**E**) BMDMs were cultured for 24 hours in custom-formulated RPMI 1640 media containing all amino acids at standard concentrations (Control) or lacking a single amino acid (Table S1), as indicated. After 24 hours, cells were primed with LPS (100 ng/mL) for 4 hours and stimulated with nigericin (5 µM) for 2 hours. Supernatants were analyzed for (**B**) IL-1β, (**C**) IL-6, and (**D**) TNF release by ELISA, and (**E**) LDH release was quantified as a measure of cytotoxicity. (B to E) Data represent the mean values of biological triplicates pooled from 3 independent experiments +/- SD. (B to D) Cytokine values from amino acid dropout conditions were normalized to control values (set at 100%) and expressed as percent change. (E) LDH release was expressed as percent cytotoxicity relative to control samples, where untreated BMDMs and detergent-permeabilized BMDMs cultured in control medium were defined as 0% and 100% cell death, respectively. AA=amino acid. OHP=hydroxyproline.

This amino acid restriction screen revealed that NLRP3-dependent and -independent inflammatory cytokine production is selectively and differentially sensitive to individual amino acids, with IL-1β release exhibiting the most dynamic regulation in both direction and magnitude (Fig. 1B). Whereas the restriction of amino acids like proline and tryptophan enhanced NLRP3-driven IL-1β release and cell death, others like glutamine, methionine, valine, and isoleucine profoundly inhibited IL-1β release and cell lysis (Fig. 1, B and E). Moreover, this screen also identified known regulators of inflammasome function and IL-1β release, such as serine (*49*) (Fig. 1, B and E). Similarly, we found that IL-6 production also displayed amino acid-specific sensitivity, and that many amino acids (e.g., methionine and serine) exerted differential effects on IL-1β release and IL-6 secretion (Fig. 1C). In contrast, tumor necrosis factor (TNF) production was minimally affected by any single amino acid, suggesting that TLR4-driven TNF secretion is largely insensitive to environmental amino acid availability (Fig. 1D). Taken together, these results indicate that discrete, non-redundant aspects of the macrophage inflammatory program are selectively regulated by environmental amino acids.

### Environmental BCAAs Control the Rate of NLRP3 Inflammasome Assembly and IL-1β Release

Among the metabolites we identified as regulators of macrophage inflammatory function, the three BCAAs—leucine, isoleucine, and valine—emerged as key drivers of NLRP3 inflammasome-triggered cell death and IL-1β production (Fig. 1, B and E). The BCAAs, characterized by their unique chemical structure and shared catabolic pathway, are essential metabolites that play central roles in regulating cell signaling, mitochondrial metabolism, transcription, and translation (*50–55*). Despite their well-established importance as physiological regulators of metabolic function (*56–60*), their role(s) in modulating critical arms of the innate immune response remain poorly understood.

To investigate how environmental BCAA availability regulates NLRP3 inflammasome activation, we cultured BMDMs in a BCAA-free medium. We first confirmed that intracellular BCAA pools are sensitive to environmental availability, finding substantially reduced intracellular leucine, isoleucine, and valine levels in macrophages cultured under BCAA-depleted conditions (fig. S1E). Consistent with our nutrient-deprivation screen, prolonged BCAA restriction inhibited NLRP3 inflammasome-driven IL-1β release and cell death (Fig. 2, A and B). In contrast, limiting the BCAAs immediately prior to LPS priming significantly impaired IL-1β release but not cell lysis (fig. S1, F and G), in keeping with our total amino acid restriction studies (fig. S1, B and C). Likewise, acutely restoring environmental BCAAs to BCAA-starved cells immediately before LPS priming rescued NLRP3-triggered IL-1β release but not cell lysis, indicating that BCAAs regulate these two processes through distinct mechanisms (fig. S1, H and I). Prolonged BCAA restriction was also found to suppress TLR4-driven IL-6 release, whereas TNF secretion was only impaired after acute BCAA withdrawal from the macrophage cell culture milieu (fig. S1, J and K). Collectively, these results reveal that BCAAs regulate macrophage inflammation in a cytokine-specific manner and that their environmental restriction impedes NLRP3 inflammasome activation.

**Fig. 2.**
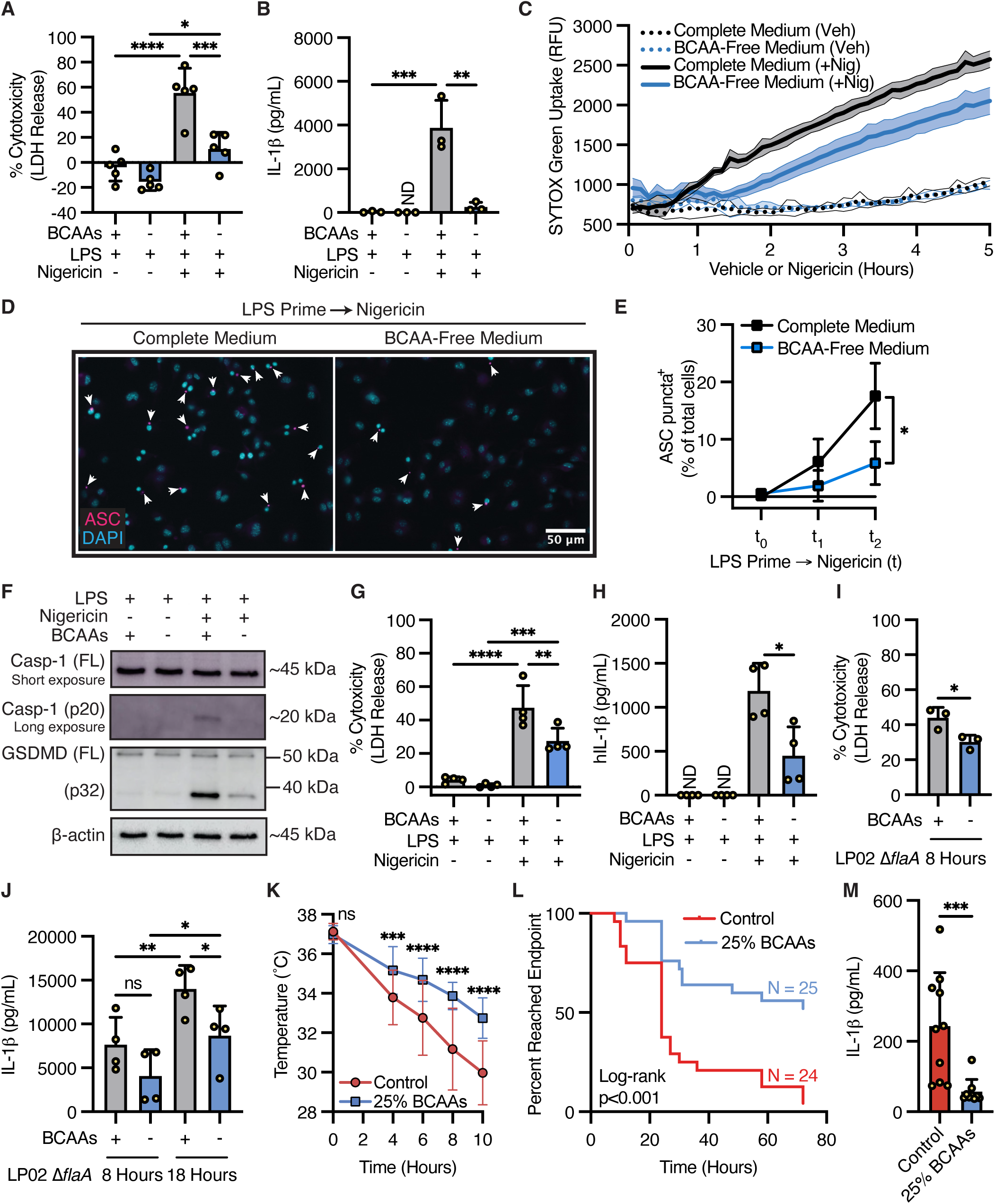
BCAAs promote inflammasome activation and IL-1β release *in vitro* and *in vivo*. (**A**) to (**F**) BMDMs were cultured for 20-24 hours in BCAA-free RPMI 1640 medium supplemented with or without leucine, isoleucine, and valine at standard concentrations before priming with LPS (100 ng/mL) for 4 hours. (**A**) LDH and (**B**) IL-1β release into supernatants was quantified by cytotoxicity assay and ELISA, respectively, after subsequent treatment of BMDMs with nigericin (5-10 µM) or vehicle control for 50-120 minutes. (**C**) SYTOX dye uptake was measured as relative fluorescence units (RFU) over 5 hours in LPS-primed BMDMs treated with 5 µM nigericin (+Nig) or vehicle control (Veh). (**D**) ASC puncta were imaged in ASC-citrine BMDMs by fluorescence microscopy after nigericin stimulation (5 µM, 40 minutes) and (**E**) quantified at 3 timepoints, where t_0_=0 minutes, t_1_=20-30 minutes, and t_2_=40-50 minutes after nigericin treatment. (**F**) Cleavage of caspase-1 and GSDMD was assessed by Western blot of BMDM lysates after LPS priming and 45 minutes of nigericin (10 µM) or vehicle control treatment. (**G**) and (**H**) hMDMs derived from 4 independent donors were cultured for 48 hours in BCAA-free RPMI 1640 medium supplemented with or without leucine, isoleucine, and valine at standard concentrations before priming with LPS (100 ng/mL) for 4 hours. (**G**) LDH and (**H**) human IL-1β (hIL-1β) release into supernatants was quantified by cytotoxicity assay and ELISA, respectively, after subsequent treatment of hMDMs with nigericin (60 µM) or vehicle control for 3-5 hours. (**I**) and (**J**) BMDMs were cultured for 20-24 hours in BCAA-free RPMI 1640 medium supplemented with or without leucine, isoleucine, and valine at standard concentrations before priming with LPS (100 ng/mL) for 4 hours followed by infection with *Legionella pneumophila* (LP02 Δ*flaA*, MOI=50). (**I**) LDH release was measured by cytotoxicity assay at 8 hours post-infection, and (**J**) IL-1β release was measured by ELISA at 8 and 18 hours post-infection. (**K**) to (**M**) 6-7-week-old mice were placed on a complete (Control) amino acid diet or BCAA-restricted (25% BCAAs) diet for 5 weeks before initiating endotoxemia. (**K**) Rectal temperatures were measured and (**L**) endpoint analysis was tracked in mice after 5 mg/kg intraperitoneal LPS injection. (**M**) Peritoneal wash was collected 3 hours after 10 mg/kg intraperitoneal LPS injection, and IL-1β levels were measured by Luminex assay. Data represent (A, B, and G to J) the mean values of biological duplicates or triplicates pooled from ≥3 independent experiments +/- SD, (C) the mean values +/- SD of biological replicates (n=4) from 1 experiment representative of 3 independent experiments, (D) 1 field of view representative of 20 captured across 2 cover slips per condition from 1 experiment representative of 3 independent experiments, (E) the percentage of macrophages containing an ASC speck (>400 cells counted across 2 cover slips per condition for each experiment), with results pooled from 3 independent experiments +/- SD, (F) results from 1 experiment representative of 3 independent experiments, (K) the mean rectal temperatures +/-SD of complete chow (n=28; 24 males and 4 females) and BCAA-restricted chow (n=29; 22 males and 7 females) mice, (L) the percentage of mice on complete (n=24; 20 males and 4 females) or BCAA-restricted (n=25; 18 males and 7 females) diets that survived or did not reach the endpoint temperature over 72 hours after LPS injection, and (M) cytokine values +/- SD, where each data point represents IL-1β quantified from an individual mouse (n=10 males on a complete diet, n=9 males on a BCAA-restricted diet). Statistical significance was determined using (A, B, G, and J) two-way ANOVA, (E, H, I, and K) unpaired student’s t-test, (L) log-rank test, and (M) Mann-Whitney test. FL=full length. ns=not significant. ND=not detectable. *p < 0.05 **p < 0.01 ***p < 0.001 ****p < 0.0001

Pyroptosis is a staged process driven by inflammasome activation that is carried out over a period of time. To discern whether environmental BCAAs were absolutely required for pyroptosis or instead altered the kinetics of cell death, we measured fluorescent dye uptake from the cell culture medium—a process dependent on GSDMD pore formation (*61*). We found that prolonged BCAA deprivation markedly delayed dye uptake, indicating slowed inflammasome activation, with BCAA-starved BMDMs failing to reach dye uptake levels observed under control conditions (Fig. 2C). This delay in NLRP3-initiated cell death upon BCAA restriction corresponded with delayed formation of NLRP3 inflammasome specks (*62*) (Fig. 2, D and E). Accordingly, we observed that BCAA deprivation resulted in reduced caspase-1 and GSDMD processing upon NLRP3 stimulation (Fig. 2F). In contrast to prolonged restriction, acute BCAA restriction minimally impacted caspase-1 and GSDMD cleavage (fig. S1L). Altogether, these data demonstrate that environmental BCAA availability governs the kinetics of NLRP3 inflammasome assembly and activation.

To investigate whether environmental BCAA levels similarly regulate NLRP3 inflammasome function in human cells, we employed primary human monocyte-derived macrophages (hMDMs). Notably, 24 hours of BCAA restriction prior to LPS priming significantly reduced nigericin-triggered IL-1β release without affecting hMDM death (fig. S2, A and B). However, extending the BCAA-restriction period to 48 hours more markedly suppressed LPS-induced pro-IL-1β priming (fig. S2C) and nigericin-triggered hMDM lysis with a corresponding reduction in IL-1β release (Fig. 2, G and H). These findings suggest that both mouse and human macrophages similarly require BCAAs for full activation of the NLRP3 inflammasome but adapt to environmental BCAA deprivation with distinct kinetics.

We next aimed to investigate whether inflammasome assembly and programmed cell death were similarly impacted during infection with an intracellular pathogen and turned to *Legionella pneumophila*, for which innate immune defense requires a multifaceted inflammasome response (*63–66*). Wild-type *Legionella* rapidly triggers NLR family CARD domain-containing protein 4 (NLRC4)-mediated pyroptosis through intracellular flagellin sensing (*67, 68*). To isolate the effects of environmental BCAAs primarily on caspase-11 and NLRP3-caspase-1 activation, we used a flagellin-deficient strain, LP02 Δ*flaA*, to avoid NLRC4-dependent cell death (*69*). Further, this bacterial strain is a thymidine auxotroph (i.e., non-replicating in the absence of exogenously added thymidine), allowing us to eliminate the potential confounding effects that replicating pathogens might have on cellular amino acid availability. Consistent with our sterile NLRP3 inflammasome (Fig. 2, A and B) and caspase-11 (fig. S2, F to H) activation results, BCAA starvation significantly inhibited *Legionella*-triggered macrophage cell death and IL-1β release in both unprimed (fig. S2, D and E) and LPS-primed BMDMs (Fig. 2, I and J).

We next explored how dietary BCAA availability tunes inflammasome activity *in vivo*. To achieve this, we maintained mice on a BCAA-limited diet (*70*) for 5 weeks before subjecting animals to an LPS-induced model of endotoxic shock (fig. S2I), an inflammasome- and IL-1-driven model of acute inflammation (*71, 72*). We found that dietary BCAA restriction protected mice against endotoxemia-triggered hypothermia (Fig. 2K) and markedly improved survival relative to control-diet mice after LPS injection (Fig. 2L), along with a corresponding decrease in IL-1β levels *in vivo* (Fig. 2M). In contrast, the production of several other cytokines and chemokines (e.g., TNF, macrophage inflammatory protein-1 [MIP-1], and keratinocyte-derived chemokine [KC], among others) was unchanged in BCAA-starved mice (fig. S2J).

Collectively, these data indicate that environmental BCAAs are key regulators of NLRP3 inflammasome assembly and subsequent pyroptosis and IL-1 release. Further, our findings suggest that major effector functions of the innate immune system are regulated by dietary BCAA intake, underscoring the nutritional tunability of acute inflammation *in vivo*.

### The BCAAs Selectively Shape a Macrophage’s Transcriptional and Translational Program Driven by TLR4 Signaling

NLRP3 inflammasome activation requires two signals: a priming stimulus to induce NLRP3 and IL-1 family cytokine expression and a second PAMP and/or DAMP signal to trigger inflammasome assembly. We therefore sought to investigate how environmental BCAAs influence each of these regulatory steps. Sustained withdrawal of the BCAAs significantly impaired TLR4-induced *Il1b* but not *Nlrp3* mRNA expression in BMDMs (Fig. 3, A and B), indicating transcript-specific regulation. This selective inhibition of *Il1b* expression was also observed during acute BCAA starvation, when the induction of *Nlrp3* mRNA remained unaffected (fig. S3, A and B). These results suggest that environmental BCAAs are required to support the transcriptional priming of IL-1β but not NLRP3.

**Fig. 3.**
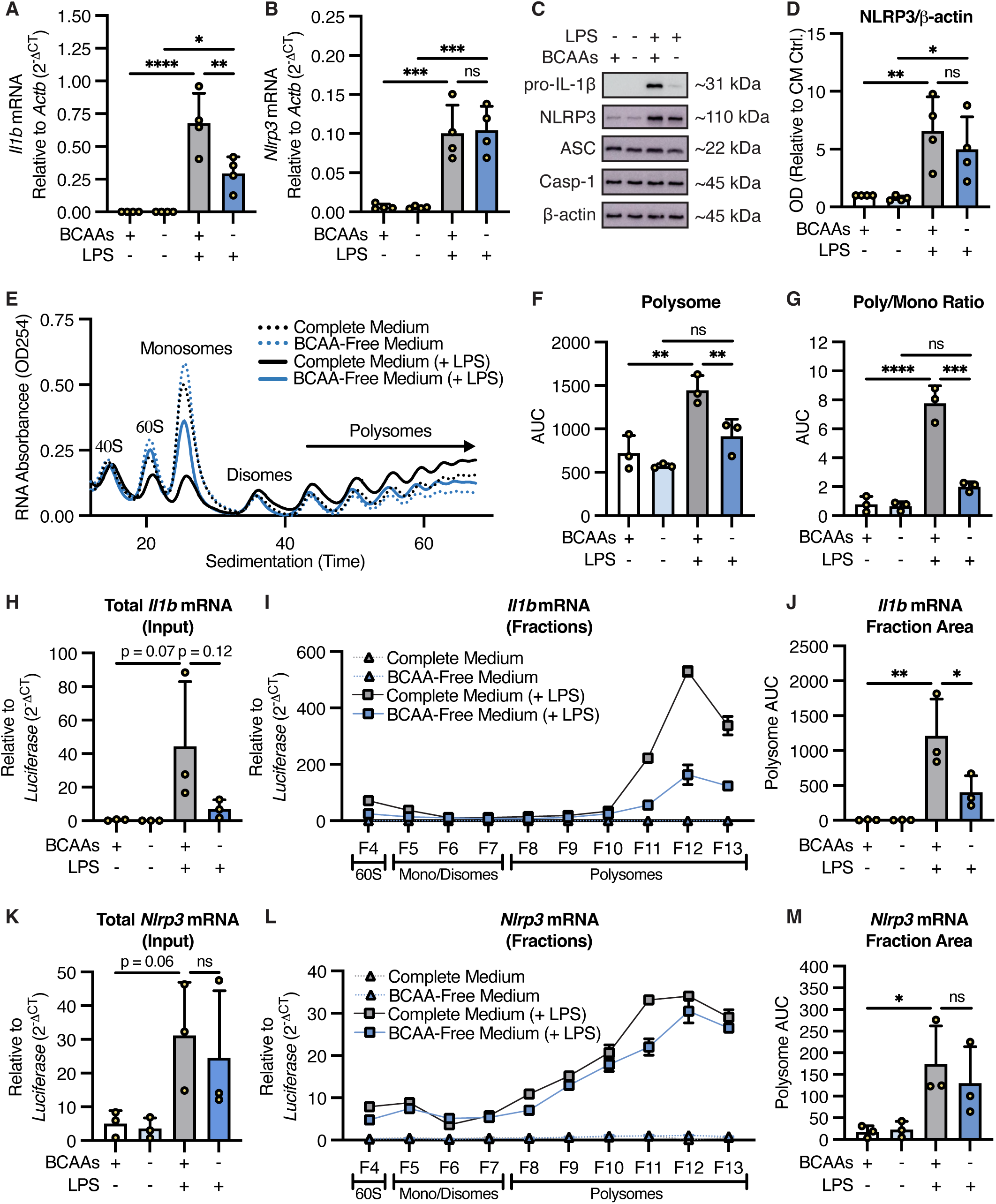
Environmental BCAAs transcriptionally and translationally license pro-IL-1β but not NLRP3 priming. (**A**) to (**M**) BMDMs were cultured for 20-24 hours in BCAA-free RPMI 1640 medium supplemented with or without leucine, isoleucine, and valine at standard concentrations before priming with LPS (100 ng/mL) or vehicle control for 4 hours. Lysates were collected for RT-qPCR analysis of (**A**) *Il1b* and (**B**) *Nlrp3* mRNA or (**C**) Western blot analysis of pro-IL-1β, NLRP3, ASC, and caspase-1 protein. (**D**) NLRP3 protein levels were measured by optical density (OD) analysis and normalized to vehicle-treated control (Ctrl.) BMDMs that were cultured in complete medium (CM). (**E**) to (**M**) Lysates were collected for polysome profiling. (**E**) Total RNA absorbance was quantified across sucrose gradient fractions with (**F**) the abundance of polysome-associated RNA and (**G**) ratio of polysome-to-monosome-associated RNA calculated by area under the curve (AUC) analysis. (**H**) *Il1b* mRNA levels were quantified in unfractionated whole-cell lysate (Input) samples and (**I**) in 60S, monosome, disome, and polysome fractions, with expression normalized to an equivalent amount of *Luciferase* mRNA spiked into each sample. (**J**) Polysome-associated *Il1b* levels were measured by AUC analysis of fractions 8 to 13 (F8-F13 in [I]). (**K**) *Nlrp3* mRNA levels were quantified in unfractionated whole-cell lysate (Input) samples and (**L**) in 60S, monosome, disome, and polysome fractions, with expression normalized to *Luciferase* mRNA. (**M**) Polysome-associated *Nlrp3* levels were measured by AUC analysis of fractions 8 to 13 (F8-F13 in [L]). Data represent (A and B) the mean values of biological duplicates pooled from 4 independent experiments +/- SD, (C and E) results from 1 experiment representative of 3 independent experiments, (D) normalized densitometry values pooled from 4 independent experiments +/- SD, (F, G, J, and M) AUC data pooled from 3 independent experiments +/- SD, (H and K) the mean values of technical triplicates pooled from 3 independent experiments +/- SD, and (I and L) RT-qPCR results showing technical triplicates +/- SD from 1 experiment representative of 3 independent experiments. (A, B, D, F to H, J, K, and M) Statistical significance was determined using two-way ANOVA. ns=not significant. *p < 0.05 **p < 0.01 ***p < 0.001 ****p < 0.0001

We next examined whether BCAA availability impacts protein expression and stability of inflammasome components. Western blot analysis of BMDM lysates showed substantial inhibition of TLR4-driven priming of pro-IL-1β, but not NLRP3, following both immediate and prolonged BCAA restriction (Fig. 3, C and D and fig. S3C). Moreover, global levels of constitutively expressed inflammasome components, ASC and caspase-1, were unaffected by acute and sustained BCAA withdrawal (Fig. 3C and fig. S3C). To distinguish between changes in mRNA and protein induction versus stability, we performed actinomycin D and cycloheximide (CHX) chase assays to measure pro-IL-1β, NLRP3, ASC, and caspase-1 mRNA and protein decay rates, respectively, in the presence and absence of environmental BCAAs. Consistent with its unique sensitivity to BCAA availability, LPS-primed *Il1b* mRNA was destabilized after prolonged BCAA deprivation, whereas *Nlrp3*, *Pycard* (i.e., ASC mRNA), and *Casp1* mRNA stability was only modestly decreased (fig. S3, D to G). We observed similar results when measuring protein decay rates. Consistent with its short half-life (*73*), pro-IL-1β showed a slight acceleration in its decay under BCAA-deprivation conditions (fig. S3H). However, ASC and caspase-1 were stably expressed in both the presence and absence of the BCAAs, and LPS-primed NLRP3 showed very modest destabilization following BCAA starvation (fig. S3H). Together, these data indicate that the NLRP3 inflammasome’s sensitivity to environmental BCAAs is not due to major changes in the initiation and/or stabilization of NLRP3 priming. However, our findings reveal specificity in the biosynthetic program shaped by the BCAAs, with TLR4-induced IL-1β mRNA and protein levels being particularly dependent on environmental BCAA availability.

To better resolve the specific stage of gene regulation that environmental BCAAs control inflammasome assembly, we next performed polysome profiling. Relative to resting BMDMs, LPS stimulation resulted in a global loss of transcripts associated with monosomes and a corresponding increase in transcripts associated with polysomes, consistent with an overall increase in protein translation (Fig. 3, E and F and fig. S4A). This TLR4-driven global translational bursting required environmental BCAAs, with BCAA-restricted BMDMs displaying modest changes in their pools of monosome- and polysome-associated RNAs relative to LPS-stimulated BMDMs cultured under control metabolic conditions (Fig. 3, E and F and fig. S4A). These differences were further reflected in the polysome-to-monosome ratio in LPS-treated BMDMs, with BCAA restriction significantly impairing increased translational capacity (Fig. 3G). Corroborating these data, puromycin incorporation was diminished in BCAA-restricted cells following LPS stimulation (fig. S4B). Collectively, these findings demonstrate that environmental BCAAs are required for the global increase in protein translation downstream of LPS sensing.

Next, to investigate whether these effects were truly global in nature or if specific genes were sensitive to environmental BCAAs, we purified RNA from monosome and polysome fractions and performed RT-qPCR, using unfractionated lysates (i.e., “Input”) as a control for total transcript abundance. In keeping with our conventional RT-qPCR data (Fig. 3A), we observed a loss of LPS-induced *Il1b* transcriptional priming in input samples collected from BCAA-starved BMDMs relative to those cultured under control metabolic conditions (Fig. 3H). This loss of *Il1b* in unfractionated lysates paralleled quantitatively similar decreases in LPS-driven polysome *Il1b* mRNA levels after BCAA deprivation (Fig. 3, I and J). These results suggest that environmental BCAAs regulate IL-1β transcription and, in turn, translation downstream of LPS sensing. Conversely, global mRNA expression and polysome translation of TLR4-induced *Nlrp3* was minimally affected by environmental BCAA levels (Fig. 3, K to M). Providing further evidence of gene-specific regulation, we observed a modest requirement for BCAAs in the transcriptional and translational priming of IL-1α, but not caspase-11 (fig. S4, C to J).

Altogether, these data demonstrate that while the BCAAs globally support protein translation downstream of TLR4 signaling, these changes occur in a transcript-specific manner. Moreover, we find that BCAAs regulate discrete facets of inflammasome biology. Although environmental BCAAs regulate the kinetics of NLRP3 inflammasome assembly and activation, these effects cannot be solely explained by changes in NLRP3 priming at the level of transcription or translation. However, we observed a key requirement for environmental BCAAs to support the TLR4-induced expression of IL-1β, an effect driven by both transcriptional and translational mechanisms.

### BCAA Sensing and mTOR Signaling Converge to Promote IL-1β Priming and NLRP3 Inflammasome Activation

In addition to serving as substrates for protein synthesis, the BCAAs assume myriad functions within the cell. BCAAs can be catabolized to promote oxidative ATP production, a process irreversibly driven by the branched-chain α-ketoacid dehydrogenase (BCKDH) complex localized to the mitochondrial inner membrane (*74*). BCAAs are also sensed by mTORC1, a major biosynthetic signaling hub governing anabolic growth and protein translation (*75–78*). Further, cell stress imposed by environmental amino acid restriction can trigger the integrated stress response (ISR) via general control nonderepressible 2 (GCN2), a protein kinase and sensor of amino acid starvation that regulates cellular stress adaptation at the level of transcriptional and translational reprogramming (*79–81*). Given these multifaceted arms of control (fig. S5A), we sought to systematically investigate the mechanism(s) by which environmental BCAAs license TLR4-driven pro-IL-1β priming and NLRP3 inflammasome activation.

We first tested whether BCAA oxidation via the BCKDH complex mechanistically links environmental BCAA availability to macrophage inflammasome function. We generated BMDMs from *LysM-Cre*^+/-^;*Dbt*^fl/fl^ mice containing a myeloid-specific deletion of dihydrolipoamide branched-chain transacylase (DBT), the key enzymatic component of the BCKDH complex responsible for BCAA catabolism (*82*), resulting in successful ablation of DBT within BMDMs (fig. S5B). We found that DBT-deficient BMDMs were as capable of priming and releasing IL-1β following NLRP3 activation as were wild-type BMDMs (fig. S5, B and C). Wild-type and DBT-deficient BMDMs also exhibited similar levels of cell death after LPS priming and nigericin treatment (fig. S5D). Finally, wild-type and DBT-deficient BMDMs were equally sensitive to the inhibitory effects of BCAA starvation on TLR4-driven pro-IL-1β priming and NLRP3-triggered cell death and IL-1β release (fig. S5, B to D). These data indicate that BCAA catabolism downstream of the BCKDH complex does not regulate NLRP3 inflammasome activation in macrophages.

Next, we investigated whether activation of the ISR in macrophages contributes to the suppressive effects of BCAA deprivation on NLRP3 inflammasome activation and IL-1β priming and release. We first measured the expression of ATF4 (activating transcription factor 4), a transcriptional activator of stress adaptation whose translation is initiated upon ISR engagement (*81, 83*). Consistent with the literature, we observed enhanced ATF4 protein levels as well as *Atf4* mRNA polysome association in BMDMs upon environmental BCAA restriction in LPS-stimulated cells, indicating robust activation of the ISR (fig. S5, E to H). Even so, treatment with the GCN2 inhibitor GCN2iB failed to negate the suppressive effects of BCAA deprivation on TLR4-induced pro-IL-1β priming and NLRP3-triggered cell death, despite blocking ATF4 induction in BCAA-starved BMDMs (fig. S5, I and J). To corroborate this pharmacologic approach, we generated BMDMs from GCN2-deficient mice, which similarly exhibited BCAA sensitivity to TLR4-driven pro-IL-1β priming and NLRP3-dependent cell death and IL-1β release (fig. S5, K to M). These data suggest that GCN2 leading to ISR engagement is not the primary mechanism by which environmental BCAAs regulate macrophage NLRP3-triggered pyroptosis.

These results next led us to test the role of mTOR signaling in mediating the BCAA-dependent licensing of NLRP3 inflammasome activation. We subjected BMDMs to prolonged BCAA restriction and/or pharmacologically inhibited mTOR signaling using either the allosteric mTORC1 inhibitor rapamycin or Torin 1, an ATP-competitive inhibitor of mTOR. Prolonged rapamycin treatment in a complete metabolic environment suppressed NLRP3 inflammasome-driven cell death and IL-1β release (Fig. 4, A and C). However, rapamycin treatment during BCAA starvation did not further enhance the inhibitory effect of BCAA restriction alone on NLRP3 inflammasome activation and IL-1β production (Fig. 4, A and C). These data suggest an epistatic relationship between BCAA sensing and mTORC1 signaling in promoting NLRP3 inflammasome activation. Similarly, Torin 1 dose-dependently suppressed NLRP3-driven cell death and IL-1β release in the presence, but not absence, of BCAAs (Fig. 4, B and D). We confirmed that both pharmacologic inhibition of mTOR and environmental BCAA restriction suppress LPS-driven mTOR signaling in BMDMs, as indicated by reduced phosphorylation of ribosomal protein S6 (RPS6), a direct substrate of mTOR-activated ribosomal protein S6 kinase (*77*) (Fig. 4E). Likewise, LPS-induced pro-IL-1β levels were dose-dependently inhibited by prolonged Torin 1 treatment, and this did not correspond with any notable defects in LPS-induced NLRP3 expression or global levels of ASC or caspase-1 (Fig. 4E). Nevertheless, prolonged mTOR inhibition dose-dependently suppressed NLRP3-triggered caspase-1 cleavage (Fig. 4F). These data suggest that BCAA sensing and mTOR signaling control NLRP3 inflammasome activation independently of LPS-driven NLRP3 expression. Whereas mTOR signaling was previously shown to promote ROS-dependent GSDMD oligomerization and membrane pore formation downstream of caspase-1 signaling (*43*), our data suggest that sustained amino acid restriction and mTOR inhibition restrains caspase-1 activation.

**Fig. 4.**
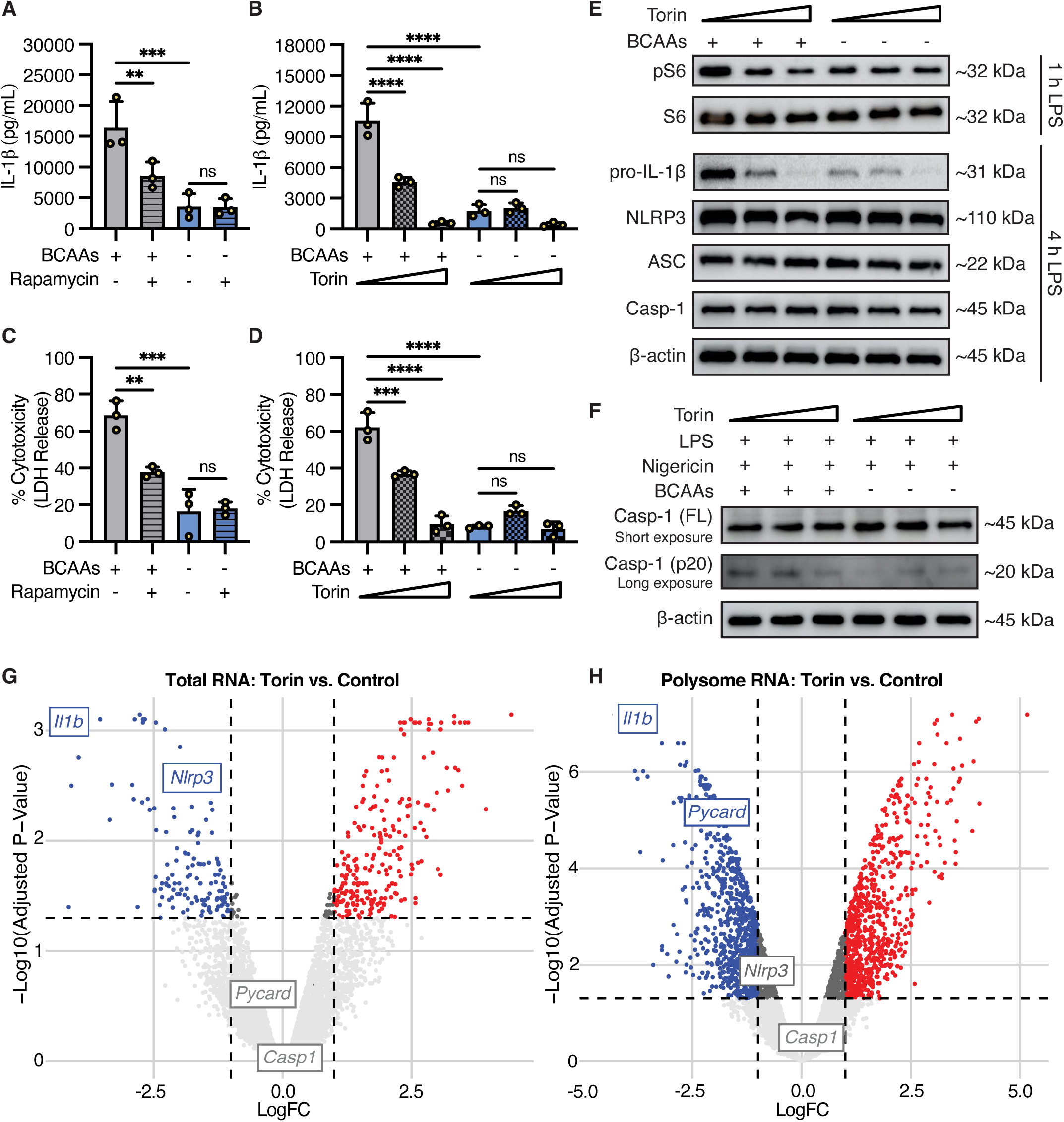
mTOR shapes a TLR4-driven biosynthetic program to promote NLRP3 inflammasome activation and IL-1β release. (**A**) to (**F**) BMDMs were cultured for 20-24 hours in BCAA-free RPMI 1640 medium supplemented with or without leucine, isoleucine, and valine at standard concentrations. (**A**) and (**C**) Rapamycin (50 nM), (**B**) and (**D**) to (**F**) Torin 1 (10 nM or 50 nM), or vehicle control was added to BMDMs for 20-24 hours parallel to BCAA starvation before priming with LPS (100 ng/mL). (**A**) and (**B**) IL-1β and (**C**) and (**D**) LDH release into supernatants was quantified by ELISA and cytotoxicity assay, respectively, after priming BMDMs with LPS for 4 hours followed by stimulation with nigericin (5 µM) for 40-140 minutes. (**E**) Phosphorylated S6 (pS6) and total S6 protein levels were assessed by Western blot of BMDM lysates 1 hour (h) after LPS stimulation, and pro-IL-1β, NLRP3, ASC, and caspase-1 protein levels were assessed 4 hours after LPS simulation. (**F**) Caspase-1 cleavage was assessed by Western blot of BMDM lysates after LPS priming (4 hours) and 45 minutes of nigericin (5 µM) treatment. (**G**) and (**H**) BMDMs were pre-treated with Torin 1 (1 µM) or left untreated (Control) for 30 minutes prior to LPS priming (100 ng/mL, 4 hours), and lysates were collected for polysome profiling. RNA sequencing was performed on (**G**) unfractionated whole-cell lysate samples or (**H**) pooled polysome fractions. Data represent (A to D) the mean values of biological triplicates pooled from 3 independent experiments +/-SD, (E and F) results from 1 experiment representative of 3 independent experiments, and (G and H) values averaged from analysis of samples collected across 3 independent experiments. (A to D) Statistical significance was determined using two-way ANOVA. FL=full length. LogFC=log fold change. ns=not significant. **p < 0.01 ***p < 0.001 ****p < 0.0001

Given this role for mTOR-dependent licensing of the NLRP3 inflammasome, we sought to more broadly characterize the biosynthetic landscape shaped by mTOR signaling during inflammasome priming. To this end, we performed polysome profiling in LPS-stimulated macrophages with or without Torin 1 treatment (fig. S6A), and we used RNA sequencing to compare total versus polysome-associated mRNA levels (fig. S6, B and C). Globally, we observed that the majority of changes following Torin 1 treatment occurred in the polysome-associated fraction, with fewer and smaller magnitude differences found in total transcript levels (fig. S6, B and C). In keeping with published observations (*84*), these findings highlight that mTOR signaling primarily controls translational rather than transcriptional rewiring during macrophage inflammatory signaling.

We next aimed to define the specific impact of mTOR activity on the transcription and translation of core NLRP3 inflammasome components. Consistent with our earlier findings, Torin 1 treatment strongly suppressed both total (Fig. 4G) and polysome-associated (Fig. 4H) *Il1b* mRNA levels during LPS stimulation. In contrast, Torin 1 treatment only modestly reduced global *Nlrp3* mRNA levels (Fig. 4G) and had no impact on *Nlrp3* abundance within polysome factions (Fig. 4H). These findings further suggest that NLRP3 translation is largely insensitive to mTOR inhibition, despite global translational suppression following Torin 1 treatment (fig. S6A).

While they are traditionally understood to be undynamic and constitutively expressed, we turned our analysis to *Casp1* and *Pycard* to ask if their translation is regulated by mTOR activity. *Casp1* mRNA levels, both total and polysome-associated, were largely unaffected by Torin 1 treatment during LPS stimulation (Fig. 4, G and H). In contrast, we observed a strong dependence on mTOR signaling for TLR4-driven ASC translation (Fig. 4, G and H). Although global *Pycard* mRNA levels remained unchanged with Torin 1 treatment during LPS stimulation (Fig. 4G), polysome-associated *Pycard* levels were substantially reduced (Fig. 4H). Therefore, mTOR signaling licenses ASC translational bursting following TLR4 stimulation.

The unexpected regulation of ASC translation downstream of mTOR next led us to ask if BCAA restriction might similarly control inflammasome assembly through an ASC-mediated mechanism. At the level of total transcript, we found that LPS stimulation led to modestly diminished *Pycard* transcript abundance in both conditions, with BCAA restriction leading to further diminished *Pycard* mRNA expression (Fig. 5A). In contrast to total transcript, we found that LPS stimulation resulted in increased *Pycard* association with polysomes, which was completely ablated in BCAA-restricted macrophages, dropping below those seen in resting control BMDMs (Fig. 5, B and C). In contrast, TLR4-driven caspase-1 transcription and translation was only modestly impaired in the absence of environmental BCAAs (fig. S6, D to F). Together, these findings demonstrate that ASC translation is dynamically engaged by TLR4 signaling in an mTOR- and BCAA-dependent manner. Our data therefore unveil a previously unidentified inflammasome priming event entailing TLR4-driven, mTOR-dependent ASC translational bursting from polyribosomes that depends on environmental BCAA availability.

**Fig. 5.**
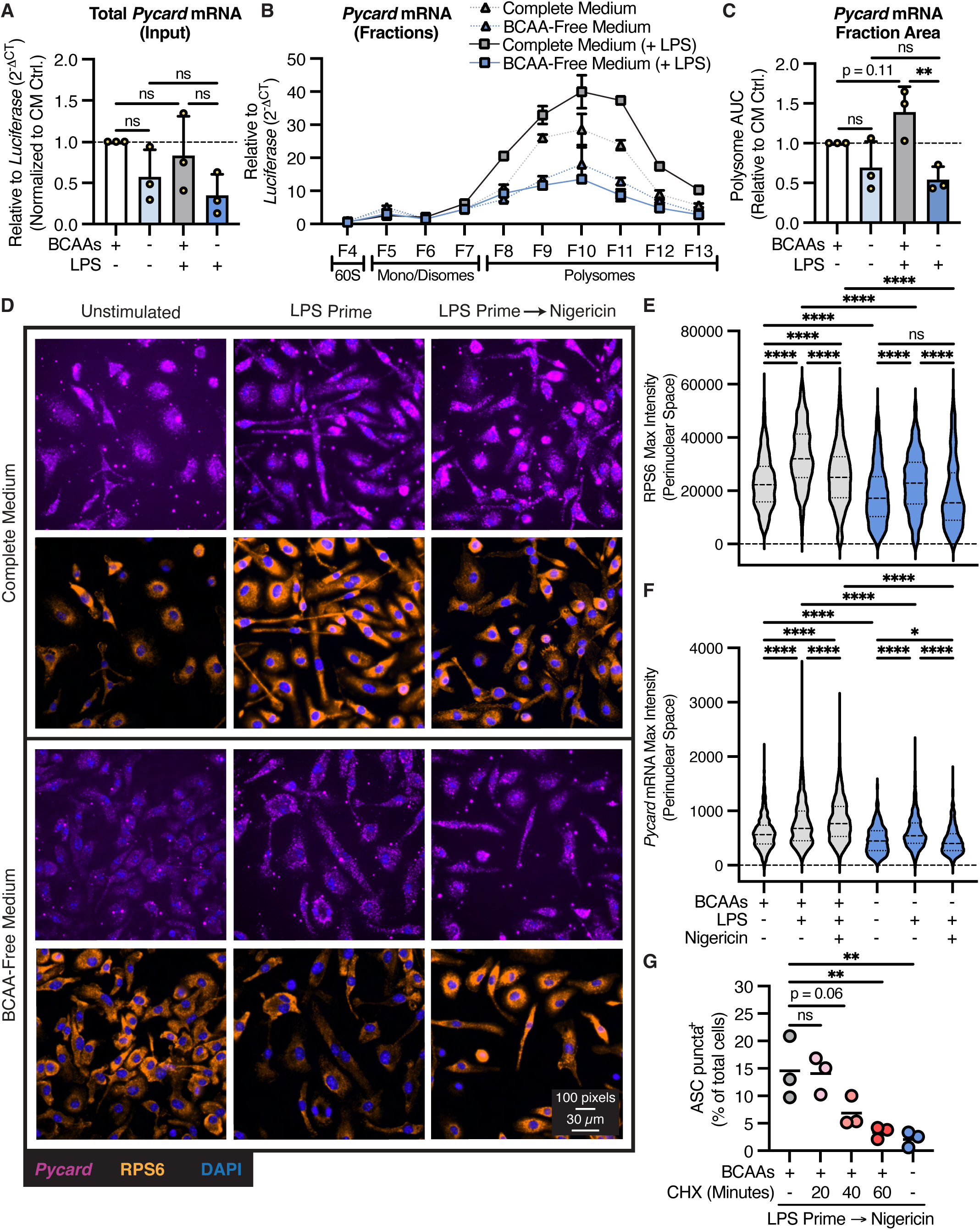
ASC translation is spatiotemporally localized to support NLRP3 inflammasome assembly. (**A**) to (**G**) BMDMs were cultured for 20-24 hours in BCAA-free RPMI 1640 medium supplemented with or without leucine, isoleucine, and valine at standard concentrations before priming with LPS (100 ng/mL) or vehicle control for 4 hours. (**A**) to (**C**) Lysates were collected for polysome profiling. (**A**) *Pycard* mRNA levels were quantified in unfractionated whole-cell lysate (Input) samples and (**B**) in 60S, monosome, disome, and polysome fractions, with expression normalized to an equivalent amount of *Luciferase* mRNA spiked into each sample. (**C**) Polysome-associated *Pycard* levels were measured by area under the curve (AUC) analysis of fractions 8 to 13 (F8-F13 in [B]) and normalized to vehicle-treated control (Ctrl.) BMDMs that were cultured in complete medium (CM). (**D**) to (**F**) Untreated, LPS-primed, and LPS-primed plus nigericin-stimulated (5 µM, 20-30 minutes) ASC-citrine BMDMs were fixed for immunofluorescence and RNA-FISH labeling of RPS6 and *Pycard* mRNA, respectively. (**D**) A representative field of view with quantification of maximum (**E**) RPS6 and (**F**) *Pycard* mRNA fluorescence intensity in a 10-pixel ring around the nucleus (Perinuclear Space). (**G**) For the final 20, 40, or 60 minutes of LPS priming, cycloheximide (CHX, 50 µg/mL) was added to halt protein translation in ASC-citrine BMDMs. The frequency of cells containing a fluorescent ASC punctum after subsequent nigericin treatment (5 µM, 30-45 minutes) was quantified by fluorescence microscopy. Data represent (A) the mean values of technical triplicates pooled from 3 independent experiments +/- SD, (B) RT-qPCR results showing technical triplicates +/- SD from 1 experiment representative of 3 independent experiments, (C) normalized AUC data pooled from 3 independent experiments +/- SD, (D) 1 field of view captured from 1 experiment representative of 2 independent experiments, (E and F) violin plots showing the median, interquartile range, and full range of the data collected from 2 independent experiments (>595 nuclei analyzed in total for each condition), and (G) the percentage of macrophages containing an ASC speck (>450 cells counted per condition for each experiment), with results pooled from 3 independent experiments. Statistical significance was determined using (A, C, E, and F) two-way ANOVA and (G) one-way ANOVA. ns=not significant. *p < 0.05 **p < 0.01 ****p < 0.0001

### Spatially Localized ASC Expression Parallels a Requirement for Protein Translation to License Assembly of the NLRP3 Inflammasome

We next asked whether this dynamic translational regulation of ASC contributes to the spatiotemporal seeding of inflammasome speck assembly. The oligomerization of ASC monomers occurs at the perinuclear space within minutes of NLRP3 activation (*33, 34*), suggesting a critical need for high local concentrations of ASC protein to enable such rapid filamentation. Moreover, recent work suggests that ASC filamentation occurs in contact with a robust ribosome network (*85*). These findings led us to hypothesize that the rapid engagement of ASC translation upon LPS stimulation might locally seed and accelerate ASC filamentation required for inflammasome assembly, and that this activity is sensitive to environmental BCAA availability.

To test this, we first performed immunofluorescence labeling of ribosomes via RPS6 staining coupled with RNA-FISH and visualized the dynamics of ribosome and ASC mRNA (*Pycard*) localization during inflammasome priming and activation. We observed that TLR4 activation yielded robust enhancement of cellular RPS6 staining (fig. S7A), an effect that extended into the perinuclear space (Fig. 5, D and E), which is classically regarded as a subcellular translation “hot spot” (*86*). These TLR4-driven increases in ribosome abundance paralleled an intensification of cellular *Pycard* mRNA staining that also localized to the perinuclear space (Fig. 5, D and F and fig. S7, B to D). These data showing concomitant increases in perinuclear RPS6 and *Pycard* mRNA localization support a model in which localized ASC translational bursting occurs during inflammasome priming. Accordingly, nigericin treatment further enhanced perinuclear *Pycard* mRNA levels, while yielding a modest reduction in RPS6 abundance (Fig. 5, D to F and fig. S7, A to D). We next tested the impact of BCAA restriction on these phenomena and observed that BCAA-starved BMDMs exhibited reduced perinuclear localization of both RPS6 and *Pycard* mRNA during NLRP3 inflammasome priming and activation (Fig. 5, D to F and fig. S7, A to D). Altogether, we demonstrate that environmental BCAA availability controls *Pycard* perinuclear localization, ASC translation, and ASC speck assembly.

In light of these findings, we next investigated whether de novo protein synthesis is required for ASC puncta formation after NLRP3 priming has already occurred. To test this, we primed BMDMs with LPS for 4 hours but added CHX to halt global translation shortly before nigericin treatment. Despite LPS-primed BMDMs maintaining robust expression of NLRP3, ASC, and caspase-1 for an hour post-CHX treatment (fig. S7E), these cells nevertheless exhibited profoundly impeded ASC speck formation (Fig. 5G). These data indicate that inflammasome assembly requires de novo translation, independent of NLRP3 priming. Collectively, our work supports a model in which environmental amino acid availability and sensing license perinuclear *Pycard* transcript localization, enabling rapid ASC translational bursting to support inflammasome assembly. This suggests that the canonical two-step model of NLRP3 inflammasome activation is insufficient. Moreover, these findings highlight how gene products otherwise found in steady-state equilibrium like ASC can be spatially and kinetically regulated to control critical cellular processes.

## DISCUSSION

Pathogen sensing initiates a metabolically intensive host defense program that requires tight coordination between environmental nutrient sensing and consumption, intracellular metabolic signaling, bioenergetic regulation, and macromolecule synthesis (*87–89*). Given their roles as both anabolic signaling molecules and substrates for ATP production and protein synthesis, amino acids control key metabolic checkpoints that govern transcriptional and translational cellular processes required for host defense. Yet despite this, the specific roles that discrete amino acids play in regulating major effector arms of the innate immune response have remained poorly defined.

Here, we demonstrate that environmental BCAAs profoundly regulate NLRP3 inflammasome activation. Specifically, our data show that BCAA sensing licenses the transcriptional and translational programs required for TLR4-driven IL-1β priming and NLRP3 inflammasome activation through mTOR-dependent mechanisms. Most notably, we identify a previously unrecognized layer of translational control governing ASC synthesis, revealing that ASC undergoes BCAA- and mTOR-dependent translational bursting during TLR4-induced inflammasome priming. These findings challenge the prevailing view of ASC as a constitutively expressed, biosynthetically static inflammasome component and instead establish it as a dynamically regulated protein whose expression is acutely controlled at the level of translation.

Mechanistically, our data suggest that TLR4 activation initiates a spatially organized translational program that concentrates ribosomes and ASC mRNA in the perinuclear region, the canonical site of ASC speck formation (*33, 34*). This coordinated localization is lost under BCAA starvation when ASC translation is impaired. Moreover, we find that transient blockade of protein synthesis—even after achieving stable expression of NLRP3—disrupts macrophage assembly of the ASC speck. These data collectively suggest a functional requirement for NLRP3-independent protein synthesis to promote inflammasome assembly and support a model in which localized ASC translation acts as a key rate-limiting step for ASC speck formation.

Why might the bioenergetics of inflammasome assembly favor seeding by a small but spatially concentrated pool of newly synthesized ASC, rather than relying on pre-existing protein that is diffusely distributed throughout the cell? Although ASC is maintained at supersaturated concentrations sufficient for spontaneous assembly, an intrinsic nucleation barrier prevents stochastic filament formation under resting conditions (*90*). A spatially concentrated burst of ASC translation may more efficiently overcome this barrier by generating a local pool of nascent ASC that acts as a nucleation “seed,” thereby tipping the cell toward rapid and robust filament polymerization. Consistent with this model, BCAA starvation delays but does not completely block ASC speck formation, suggesting that the pre-existing ASC pool provides a foundation for inflammasome assembly, while new ASC synthesis helps accelerate filamentation to more efficiently cross the critical nucleation threshold required for inflammasome activation (summarized in fig. S7F).

Although our data support various elements of this model, several important questions remain unanswered and will require more rigorous investigation. Most notably, how is ASC translation selectively upregulated and spatially confined to the perinuclear space during inflammasome priming and assembly? Addressing this question will be an important next step toward understanding how inflammasome assembly is spatiotemporally regulated.

In addition to its role in regulating ASC translation, our data implicate mTOR signaling as a critical transcriptional and translational driver of IL-1β priming. The strong requirement for mTOR to license TLR4-induced IL-1β transcription was notable given our data suggesting translation as the predominant means of regulation exerted by mTOR signaling in macrophages. How mTOR shapes a cell’s transcriptional landscape is poorly understood, but the extant body of literature suggests that mTOR controls gene transcription by regulating the expression and/or activity of key transcription factors (*91–93*). Among these, synthesis of hypoxia-inducible factor 1α (HIF-1α), a master regulator of energy metabolism, is translationally driven by mTOR (*94*). Further, HIF-1α undergoes dynamic regulation during TLR4 activation, when it translocates from the cytosol to the nucleus and selectively binds to the *Il1b* promoter region to transcriptionally drive its expression (*87*). However, whether HIF-1α, or any other mTOR-sensitive transcription factor, directly links nutrient sensing to IL-1 gene regulation is unknown, and we are actively exploring this gap in knowledge.

Another intriguing and yet unresolved observation pertains to the kinetics by which macrophages adapt to a BCAA-starved environment. We find that limiting amino acids does not immediately suppress pyroptotic signaling but instead requires a prolonged period of nutrient restriction to functionally impair NLRP3 inflammasome activation. This temporal delay suggests that macrophages possess a short-term capacity to buffer against acute fluctuations in amino acid availability, potentially by mobilizing intracellular amino acid stores or recycling extracellular proteins. However, with sustained amino acid stress, more fundamental reprogramming of host cell machinery may occur and will require further mechanistic investigation.

Lastly, our *in vivo* data demonstrating that dietary BCAA restriction protects mice from endotoxic shock highlight a broader physiological relevance for BCAA sensing in the regulation of inflammasome-driven inflammation. Several chronic metabolic diseases and syndromes, including obesity, type 2 diabetes, cardiovascular disease, and aging, feature elevated circulating BCAA levels (*56–60, 95*) and enhanced NLRP3 inflammasome activation (*15, 20, 21*), yet the mechanistic connections between these observations remain unresolved. Our findings raise the possibility that excess BCAAs serve not only as biomarkers of metabolic dysfunction but also as metabolic signals that license inflammasome function. In principle, sustained BCAA abundance could augment the priming arm of the inflammasome by chronically enhancing mTOR-dependent transcriptional and translational programs, thereby lowering the threshold for ASC nucleation and IL-1β production. This scenario suggests that cells in BCAA-rich environments might exist in “hyperprimed” inflammasome states, rendering them more responsive to inflammatory cues like metabolic damage signals that prompt NLRP3 activation (*15*). Although speculative, this model provides a potential mechanistic framework for understanding how nutrient excess in metabolic disease may interface with innate immune signaling to drive chronic, low-grade inflammation. Our dietary restriction data thus raise the intriguing possibility that manipulating systemic BCAA availability could modulate inflammasome activity in metabolic disease contexts, providing an immunometabolic entry point for therapeutically tuning pathological NLRP3 activation.

In summary, our study uncovers a critical role for environmental BCAAs and mTOR signaling in regulating key biosynthetic checkpoints that control inflammasome activation in macrophages. We identify ASC translation as a dynamically regulated, nutrient-sensitive, and spatially organized process that potentially operates as a rate-limiting step for inflammasome assembly, challenging the longstanding view of ASC as a biosynthetically static structural component. Moreover, our data position the amino acid environment and mTOR signaling as key transcriptional regulators of IL-1β. Together, these findings reveal that nutrient availability is not merely permissive for immune activation but actively shapes the timing, localization, and efficiency of inflammasome responses through coordinated transcriptional and translational programs.

## MATERIALS AND METHODS

### Mice

All mice were maintained under specific pathogen-free conditions at the University of Pennsylvania or Children’s Hospital of Philadelphia. C57BL/6J and B6.Cg-*Gt(ROSA)26Sor^tm1.1(CAG-Pycard/mCitrine*,-CD2*)Dtg^*/J mice (wild-type and ASC-citrine reporter mice, respectively) were either purchased from Jackson Laboratories (#000664 and #030744) or bred in house. Bone marrow from B6.129S6-*Eif2ak4^tm1.2Dron^*/J (GCN2 knockout) and corresponding wild-type (age-, sex-, and facility-matched C57BL/6J) mice was kindly provided by Dr. Mary O’Riordan. For DBT knockout experiments, B6.129P2-*Lyz2^tm1(cre)Ifo^*/J (*LysM-Cre*) mice were purchased from Jackson Laboratories (#004781) and crossed to *Dbt*^fl/fl^ mice kindly provided by Dr. Rebecca Ahrens-Nicklas to generate *LysM-Cre*^+/-^;*Dbt*^fl/fl^ mice as a source of DBT-deficient BMDMs. Bone marrow from these mice alongside age- and sex-matched *LysM-Cre*^+/-^ mice (used to generate DBT wild-type BMDMs) was harvested for *in vitro* experiments. All animal experiments were conducted in accordance with federal regulations outlined in the Animal Welfare Act (AWA), the National Institutes of Health’s Guide for the Care and Use of Laboratory Animals, and the policies of the University of Pennsylvania and Children’s Hospital of Philadelphia Institutional Animal Care and Use Committees (IACUC). All procedures were reviewed and approved by the University of Pennsylvania’s IACUC under protocol #804928 and Children’s Hospital of Philadelphia’s IACUC under protocol #IAC 24-001325.

### Differentiation and Culture of Primary Macrophages

To generate BMDMs, bone marrow cells were harvested from the leg bones of mice and either freshly used or frozen down for thawing and plating. 5 x 10^6^ bone marrow cells were plated per 100/20 mm petri dish in 10 mL macrophage growth medium containing RPMI 1640 (Corning #10-041-CM) supplemented with 20% fetal bovine serum (FBS, GeminiBio #100-106), 20 ng/mL recombinant murine macrophage colony-stimulating factor (M-CSF, PeproTech #315-02), and 1% penicillin/streptomycin (P/S, Gibco #15140-122). Cells were incubated at 37°C in a 5% CO_2_ incubator for 6 days to allow for differentiation with addition of 10 mL fresh growth medium per petri dish on day 4. To generate hMDMs, primary monocytes were isolated from anonymous healthy human donors by the University of Pennsylvania Human Immunology Core. 5 x 10^6^ monocytes were plated per 100/20 mm petri dish in 10 mL growth medium containing RPMI 1640 medium supplemented with 20% FBS, 20 ng/mL human M-CSF (GeminiBio #300-161P), and 1% P/S. hMDMs were also differentiated at 37°C in a 5% CO_2_ incubator for 6 days with addition of 10 mL fresh growth medium on day 3. After differentiation, macrophages were detached from plastic using cold PBS containing EDTA (2 mM EDTA for BMDMs and 10 mM EDTA for hMDMs) and re-plated in macrophage plating medium containing RPMI 1640 supplemented with 10% FBS and 10 ng/mL M-CSF for downstream experimentation using the culture media described below.

### Amino Acid-Defined Culture Media

To titrate total amino acids in cell culture, a complete RPMI 1640 medium containing 100% amino acid levels (United States Biological #R8999) was prepared according to manufacturer’s instructions and diluted to 10% using an amino acid-free RPMI 1640 medium (United States Biological #R9010-01) supplemented with standard glucose levels. These 100% and 10% amino acid media were then used for cell culture experiments.

To restrict individual amino acids in cell culture, we developed a nutrient-sensing screen guided by RPMI 1640’s standard formulation. The details regarding development of this screen are comprehensively outlined in Table S1 and briefly described here. First, using the amino acid-free RPMI 1640 medium specified above, we generated 100x stock solutions of every protein-coding amino acid represented in a commercial RPMI 1640 formulation. Then, we developed 21 distinct culture media, including a complete medium containing the full profile of amino acids and 20 dropout media, each one lacking a single amino acid. This was achieved by selectively supplementing amino acid-free RPMI 1640 with different combinations of amino acid solutions. For Fig. 1 data derived from this screen, three separate batches of media were generated using the same 100x amino acid stock solutions to complete three independent experiments.

To restrict the BCAAs in cell culture, we either (1) obtained a BCAA-free RPMI 1640 medium lacking leucine, isoleucine, and valine (United States Biological #R8999-20), prepared according to manufacturer’s instructions, or (2) independently developed BCAA-free RPMI 1640 by supplementing an amino acid-free RPMI 1640 medium (MyBioSource #MBS652918) with every amino acid except for leucine, isoleucine, and valine (amino acid manufacturers and concentrations outlined in Table S1). At the time of experimentation, 100x stock solutions of leucine, isoleucine, and valine were or were not diluted 1:100 into BCAA-free RPMI 1640 to generate a 1x “Complete Medium” or “BCAA-Free Medium,” respectively.

### Inflammasome Activation and *Legionella* Infection *In Vitro*

Following harvest, BMDMs or hMDMs were diluted to 2.5 x 10^5^ cells/mL in macrophage plating medium and seeded onto tissue culture-treated plastic plates. Cells were allowed to adhere overnight. Unless otherwise specified, experiments were conducted in 24-well or 48-well plates, with each well receiving 500 µL or 250 µL of medium, corresponding to 1.25 x 10^5^ or 6.25 x 10^4^ cells per well, respectively. After adherence, macrophages were washed with amino acid-free RPMI for amino acid titration and single amino acid restriction experiments, or with BCAA-free RPMI for BCAA restriction experiments. Cells were then replenished with custom media as described above, supplemented with 10% dialyzed FBS (Cytiva #SH30079.03, lot #AF2941643) and 10 ng/mL M-CSF.

For immediate amino acid restriction (“Immediate Restriction”), macrophages were first cultured for 20-24 hours in complete medium containing 10% dialyzed FBS and 10 ng/mL M-CSF. Cells were then washed and replenished with the specified amino acid-restricted medium (containing 10% dialyzed FBS) and immediately stimulated with LPS, as described below. For prolonged amino acid restriction (“Prolonged Restriction”), macrophages were cultured for 20-24 hours in amino acid-restricted medium containing 10% dialyzed FBS and 10 ng/mL M-CSF. Cells were then replenished with fresh, identically restricted medium (again containing 10% dialyzed FBS) and stimulated with LPS. For experiments in which hMDMs were starved of the BCAAs for 48 hours, fresh medium was added to cells at 24 and 48 hours after beginning restriction.

Following amino acid deprivation, the NLRP3 inflammasome was primed and activated by sequential treatment of macrophages with 100 ng/mL LPS from *Escherichia coli* O111:B4 (Sigma-Aldrich #L2630) for 4 hours, followed by nigericin treatment. Nigericin was obtained from either EMD Millipore (#481990) or Sigma-Aldrich (#SML1779), with concentrations and treatment durations specified in the corresponding figure legends.

To activate the caspase-11 inflammasome, macrophages were first primed with 100 ng/mL LPS from *Escherichia coli* O111:B4 for 6 hours, then washed with Opti-MEM (Gibco #31985062) and transfected with 1 µg/mL LPS from *Salmonella enterica* serotype Minnesota (Sigma-Aldrich #L6261). LPS was packaged into stable complexes using FuGENE HD (Promega #E2311) and delivered to cells in Opti-MEM, as previously described (*96*). Cells were incubated with the transfection mixture for 5 hours. Flagellin-deficient *Legionella pneumophila* on a *thyA* thymidine auxotrophic background (LP02 Δ*flaA*) (*69*) was cultured on charcoal yeast extract agar (CYE) containing streptomycin and supplemented with thymidine for 4 days. Single colonies were isolated and cultured on CYE plates for 48 hours and then suspended in sterile PBS immediately prior to use. For infection experiments *in vitro*, BMDMs were either rested or primed with LPS from *Escherichia coli* O111:B4 (100 ng/mL) for 4 hours followed by infection with LP02 Δ*flaA* at a multiplicity of infection of 50 for either 8 or 18 hours prior to harvesting samples for analysis.

For chemical inhibitor studies, macrophages were treated for the time durations and using the concentrations specified in each figure legend. The following inhibitors were used: Torin 1 (Cayman Chemical #10997), rapamycin (MedChemExpress #HY-10219), GCN2iB (MedChemExpress #HY-112654), cycloheximide (Sigma-Aldrich #C4859), actinomycin D (Sigma-Aldrich #SBR00013), and Z-VAD-FMK (SM Biochemicals #SMFMK001).

### LDH Release and SYTOX Uptake Cytotoxicity Assays

At the specified timepoints after infection or inflammasome activation, tissue culture plates were centrifuged for 3 minutes at 1,200 RPM to spin down floating cells. Supernatant samples were then harvested, and LDH activity was measured by colorimetric assay (Sigma-Aldrich #11644793001), per the manufacturer’s instructions, as a readout for cell lysis. Unless otherwise noted, % cytotoxicity was calculated relative to untreated and detergent-permeabilized (1% triton x-100) wild-type BMDMs cultured in complete medium, which were set at 0% and 100% cytotoxicity, respectively.

To kinetically measure plasma membrane pore formation after inflammasome activation, cellular uptake of the SYTOX green nucleic acid stain (ThermoFisher Scientific #S7020) was quantified. To achieve this, 175 µL of a 2.5 x 10^5^ cells/mL suspension was plated per well in a 96-well, black-walled, clear-bottomed tissue culture plate. After BCAA restriction, BMDMs were primed with LPS for 4 hours, as described above, and nigericin or vehicle control was added to cells in 25 µL of an equivalent medium containing SYTOX green (1 µM final concentration). Fluorescence was measured in each well every ∼7 minutes for 5 hours using an Infinite 200 Pro plate reader (Tecan #30050303), and the mean fluorescence value from 8 reads encapsulating the entirety of each well was calculated.

### Dietary Intervention and Endotoxic Shock *In Vivo*

C57BL/6J wild-type mice were bred in house at the Children’s Hospital of Philadelphia and fed on a normal chow diet until 6-7 weeks of age. Mice were then switched to amino acid-defined diets from Teklad, including a complete “Control” chow (TD.140711) or “25% BCAAs” chow (TD.150387) (*70*) for 5 weeks before performing endotoxic shock. After 5 weeks of dietary intervention, mice were either injected intraperitoneally with 5 mg/kg or 10 mg/kg LPS (Sigma-Aldrich #L2630), as specified in each figure legend.

Beginning at the time of LPS injection and ending at 10 hours post-injection, rectal temperatures were measured using a single-input thermocouple thermometer (Cole-Palmer #EW-20250-91). Additionally, mice were regularly monitored over 72 hours to track survival, with humane euthanasia performed upon detecting a rectal temperature of ≤27°C. To collect peritoneal wash for measuring cytokine levels, mice were humanely euthanized at 3 hours post-injection, and 3 mL PBS + 1% bovine serum albumin (BSA) was instilled into and then retracted from the peritoneal cavity. Peritoneal washes were centrifuged to pellet and remove cells, and 500 µL of cell-free sample was spun in a 10-kDa centrifugal filter unit (Millipore #UFC8010) for 30 minutes at 3,220 x g (4°C) to concentrate cytokine levels. Concentrated samples were then diluted 1:2 in PBS + 1% BSA and submitted for Luminex assay.

### ELISA and Luminex Assay

After collecting cell culture supernatants for the LDH cytotoxicity assay (as described above), samples were further centrifuged at 2,000 rpm for 10 minutes to remove residual cell debris. The clarified supernatants were either used immediately or stored at −80°C for long-term preservation.

ELISAs for IL-1β (BioLegend #432601), IL-6 (coating antibody: ThermoFisher Scientific #14-7061-81; detection antibody: ThermoFisher Scientific #13-7062-81; standard: BioLegend #575709), and TNF (R&D Systems #DY410) were performed according to manufacturers’ instructions. Experimental samples were diluted in ELISA assay buffer to ensure optical density readings fell within the range of the standard curve.

After collecting peritoneal wash samples from the endotoxic shock model (as described above), samples were submitted to the University of Pennsylvania Human Immunology Core for Luminex analysis. A mouse cytokine/chemokine magnetic bead panel (premixed 32-plex; MCYTMAG-70K-PX20) was used to perform the immunology multiplex assay according to the manufacturer’s instructions.

### Western Blot

Macrophages were lysed using 1x sodium dodecyl sulfate (SDS)-polyacrylamide gel electrophoresis (PAGE) sample buffer (100 µL buffer for every 1.25 x 10^5^ cells) containing 100 mM dithiothreitol (DTT). Cell lysates were boiled for 5 minutes, separated by SDS-PAGE, and transferred onto PVDF Immobilon-P membranes (Millipore #IPVH00010). Membranes were probed with antibodies against mouse IL-1β (1:500 dilution, Cell Signaling #12242), human IL-1β (1 µg/mL final concentration, R&D Systems #MAB201), mouse NLRP3 (1:1,000 dilution, Cell Signaling #15101), mouse ASC (1:500-1:1,000 dilution, Cell Signaling #67824), mouse caspase-1 (1:500 dilution, gift of Genentech [clone 4B4.2.1]), mouse caspase-11 (1:1,000 dilution, Cell Signaling #14340), mouse GSDMD (1:500, Abcam #ab209845), mouse phospho-S6 (1:5,000 dilution, Cell Signaling #2211), mouse S6 (1:5,000 dilution, Cell Signaling #2217), mouse DBT (1:200 dilution, Proteintech #12451-1-AP), mouse GCN2 (1:500 dilution, Cell Signaling #3302), mouse ATF4 (1:1,000 dilution, Cell Signaling #82093), and puromycin (1:1,000 dilution, Sigma-Aldrich #MABE343). β-actin (1:1,000 dilution, Cell Signaling #4967) was used as a loading control. Detection was performed using HRP-conjugated anti-mouse IgG (1:2,000-1:10,000 dilution, Cell Signaling #7076), anti-rabbit IgG (1:2,000-1:10,000 dilution, Cell Signaling #7074), and anti-rat IgG (1:2,000-1:10,000 dilution, Cell Signaling #7077) antibodies and SuperSignal West Femto Maximum Sensitivity Substrate (ThermoFisher Scientific #34095), used according to the manufacturer’s instructions.

### Reverse Transcription (RT)-qPCR

To harvest RNA from whole-cell lysates for conventional RT-qPCR, BMDMs were lysed and RNA was isolated using the RNeasy Plus Mini Kit (Qiagen #74136). For harvesting RNA from polysome fractions (fractionation described below), 250 µL of each sucrose fraction was used. To compare global versus polysome-associated RNA levels, thereby distinguishing transcriptional versus translational regulation, 20 µL of input (i.e., unfractionated whole-cell lysate) RNA was also used and diluted to 250 µL with molecular biology grade water before RNA purification. Then, for each sucrose fraction and input sample, 750 µL of TRIzol LS (ThermoFisher Scientific #10296028) containing 20 pg *Luciferase* control mRNA (Promega #L4561) was added and thoroughly mixed followed by addition of 1 mL 100% ethanol. Samples were then loaded onto separate columns of a Direct-zol-96 plate (Zymo Research #R2056), and RNA was purified (with DNase I treatment) according to the manufacturer’s instructions.

Following RNA purification, reverse transcription to cDNA was performed using SuperScript II Reverse Transcriptase (ThermoFisher Scientific #18064014) and oligo (dT) primers, performed according to the manufacturer’s instructions. cDNA was diluted in molecular biology grade water and quantified via RT-qPCR using PerfeCTa SYBR Green SuperMix (Quantabio #95054) and 0.5 µM forward and reverse primers on a QuantStudio 6 Flex real-time PCR system (ThermoFisher Scientific). Primers used in this study were as follows: *Il1b* forward: 5’ - GCA ACT GTT CCT GAA CTC AAC T - 3’; *Il1b* reverse: 5’ - ATC TTT TGG GGT CCG TCA ACT - 3’; *Il1a* forward: 5’ - CGA AGA CTA CAG TTC TGC CAT T - 3’; *Il1a* reverse: 5’ - GAC GTT TCA GAG GTT CTC AGA G - 3’; *Il18* forward: 5’ - TGA AGT AAG AGG ACT GGC TGT GAC - 3’; *Il18* reverse: 5’ - ATC TTG TTG TGT CCT GGA ACA CG - 3’; *Il6* forward: 5’ - TAC CAC TTC ACA AGT CGG AGG C - 3’; *Il6* reverse: 5’ - CTG CAA GTG CAT CAT CGT TGT TC - 3’; *Tnf* forward: 5’ - CTG AAC TTC GGG GTG ATC GG - 3’; *Tnf* reverse: 5’ - GGC TTG TCA CTC GAA TTT TGA GA - 3’; *Nlrp3* forward: 5’ - ATT ACC CGC CCG AGA AAG G - 3’; *Nlrp3* reverse: 5’ - TCG CAG CAA AGA TCC ACA CAG - 3’; *Pycard* forward: 5’ - CTT AGA GAC ATG GGC TTA CAG G - 3’; *Pycard* reverse: 5’ - CTC CAG GTC CAT CAC CAA GTA G - 3’; *Casp1* forward: 5’ - ACA CGT CTT GCC CTC ATT ATC T - 3’; *Casp1* reverse: 5’ - TTT CAC CTC TTT CAC CAT CTC C - 3’; *Casp11* forward: 5’ - AGC GTT GGG TTT TTG TAG ATG C - 3’; *Casp11* reverse: 5’ - CCT TGT GAA CTC TTC AGG GGA - 3’; *Actb* forward: 5’ - GGT GTG ATG GTG GGA ATG G - 3’; *Actb* reverse: 5’ - GCC CTC GTC ACC CAC ATA GGA - 3’; *Atf4* forward: 5’ - TCA GAC ACC GGC AAG GAG GAT - 3’; *Atf4* reverse: 5’ - TCT GGC ATG GTT TCC AGG TCA TC - 3’; *Luciferase* forward: 5’ - GAG ATA CGC CCT GGT TCC TG - 3’; *Luciferase* reverse: 5’ - ATA AAT AAC GCG CCC AAC AC - 3’.

For each run, cycle threshold (Ct) values from 3 technical replicates were averaged and normalized to an appropriate control mRNA (specified in each figure) using the 2^-ΔCt^ method (*97*).

### Polysome Profiling and Sequencing

BMDMs were seeded onto 150/20 mm petri dishes (CELLTREAT Scientific Products #229650) at 2.5 x 10^5^ cells/mL (52 mL per plate). Following BCAA deprivation or Torin 1 pre-treatment and subsequent LPS stimulation (100 ng/mL) for 4 hours, BMDMs were washed with 5 mL cold PBS containing 50 µg/mL cycloheximide and lysed in 500 µL polysome extraction buffer (10 mM HEPES, 100 mM KCl, and 5 mM MgCl_2_ at pH=7.4 with 100 µg/mL cycloheximide and 2 mM DTT) containing 1% triton x-100 and 40 units/mL SUPERase•In RNase Inhibitor (ThermoFisher Scientific #AM2696). Lysates were then incubated for 30 minutes on ice with vortexing every 5-10 minutes and centrifuged at 12,000 x g for 15 minutes at 4°C to pellet cell debris. Following centrifugation, 50 µL lysate was frozen as input (i.e., unfractionated) RNA, and 400 µL lysate was loaded onto a 15-45% sucrose gradient, as described below.

Sucrose solutions were prepared in polysome extraction buffer, and gradients were prepared in SW41 ultracentrifuge tubes by mixing 15% and 45% sucrose solutions using a Biocomp Gradient Master 108. After loading lysates onto a 15-45% sucrose gradient, samples were centrifuged at 37,000 rpm for 2 hours at 4°C using the SW41 rotor of an Optima XPN-80 ultracentrifuge (Beckman Coulter Life Sciences). Gradients were then fractionated with a speed of 800 µL/minute using a Biocomp Piston Gradient Fractionator, which recorded the OD254 nm to measure RNA abundance. Fractions corresponding to one-minute intervals were collected, labeled #1-15, and stored at −80°C before RNA extraction, as described above.

For polysome RNA sequencing, RNA from sucrose gradient fractions was isolated as described above and quantified using a NanoDrop (ThermoFisher Scientific). A pooled sample containing an equal volume of RNA from polysome fractions (fractions #8-13) was generated for each sample for downstream sequencing. For total RNA sequencing, RNA from input samples was submitted in parallel.

All RNA samples were sequenced by Novogene. mRNA was purified from total RNA using poly-oligo(dT)-attached magnetic beads and subsequently fragmented. First-strand cDNA was synthesized using random hexamer primers, followed by second-strand synthesis incorporating dUTP in place of dTTP. Directional libraries were prepared through end repair, A-tailing, adapter ligation, size selection, PCR amplification, and purification. Library quality was assessed by Qubit quantification, real-time PCR, and Bioanalyzer analysis of fragment size distribution. Libraries that passed quality control were pooled according to their effective concentrations and target sequencing depth and were then subjected to Illumina sequencing.

RNA-sequencing reads were examined and filtered for quality using FastQC (v0.12.1), MultiQC (v1.25.1), and fastp (v0.22.0, using the following parameters: -q 15, -u 10, -e 15, -l 75, -y 30, --cut_front 15, --cut_tail 15, --cut_window_size 3, -g, -x, -p). Reads were then aligned to the GRCm39 mouse genome (GENCODE vM36 annotation) using STAR (v2.7.10b). The resulting alignment files were sorted and indexed with Samtools (v1.16.1). Htseq-count (v2.0.3) was used to obtain gene-level pseudocounts. EdgeR (v4.3.0) was used for downstream processing of gene counts. Genes with low expression levels were filtered with *filterByExpr*, library sizes were normalized with *calcNormFactors*, and gene-level dispersion was estimated with *estimateDisp*.

Principal component analysis was done using gene expression (normalized by logCPM) using the *prcomp* function in R. Data visualization was done with ggplot2 (v3.5.1).

Data are accessible at GEO Accession #GSE316372.

### Imaging and Quantification of Inflammasome Specks

For inflammasome speck analyses in Fig. 2, D and E, ASC-citrine BMDMs (2.5 x 10^5^ cells/mL) were seeded at 500 µL per well onto fibronectin-coated glass coverslips in 24-well plates. Coverslips were pre-coated with 500 µL of fibronectin (10 µg/mL) for 2 hours at 37°C and then rinsed twice with sterile water prior to cell seeding. Cells were cultured in amino acid-defined media and subjected to inflammasome priming followed by nigericin-triggered NLRP3 activation, as described above. At the indicated timepoints after nigericin treatment (specified in each figure legend), BMDMs were fixed by washing twice with room-temperature PBS and incubating with cold 4% paraformaldehyde on ice for 20 minutes. Coverslips were washed with PBS and mounted using DAPI Fluoromount-G (SouthernBiotech #0100-20) for imaging after drying overnight. For experiments shown in Fig. 5G, ASC-citrine BMDMs were prepared and treated as above but seeded onto 8-well chambered Permanox slides (ThermoFisher Scientific #177445) at 250 µL per well.

All samples were imaged using a Leica DM6000B upright widefield fluorescence microscope (Penn Vet Imaging Core) at 40x magnification. Citrine fluorescence was detected using GFP excitation/emission settings. Imaging parameters, including exposure time, gain, and laser intensity, were kept constant across all experimental groups.

For quantification of inflammasome specks, ASC puncta were manually counted in each field of view. The frequency of cells containing an ASC punctum was calculated by dividing the number of specks by the number of cells, as determined by automated nuclei counting in Fiji ImageJ2 (version 2.14.0/1.54f). At least 10 randomly selected fields of view per condition were analyzed across 3 independent experiments. The minimum number of cells analyzed per condition and the statistical tests used are specified in each figure legend.

### Combined Immunofluorescence and RNA-FISH

ASC-citrine BMDMs (2.5 x 10⁵ cells/mL) were seeded at 250 µL per well onto 8-well chambered Permanox slides, as described above. Cells were cultured in amino acid-defined media and either left untreated, primed with LPS, or subjected to inflammasome priming followed by NLRP3 activation with nigericin for 20-30 minutes. BMDMs were washed once with PBS and fixed in 4% formaldehyde for 10 minutes at room temperature. After fixation, cells were washed twice with PBS and permeabilized overnight in 70% ethanol at −20°C.

Combined immunofluorescence (IF) and RNA-FISH was performed using Molecular Instruments’ *HCR IF + HCR RNA-FISH protocol for mammalian cells on a chambered slide* (revision 3, 02-13-2023), following the manufacturer’s instructions without modification.

RPS6 protein was detected by IF using a rabbit primary antibody (1:200 dilution, Cell Signaling #2217) and Molecular Instruments’ HCR IF donkey anti-rabbit probe compatible with amplifier B2 (fluorophore: 546 nm). Mouse *Pycard* mRNA was detected using custom HCR RNA-FISH probes (Molecular Instruments; up to 20 binding sites across the full *Pycard* transcript) and visualized with amplifier B1 (fluorophore: 647 nm). Imaging was performed on an inverted Nikon TI-E microscope at 60x magnification across 5 Z-planes.

RNA-FISH images were analyzed using the custom imaging software NimbusImage (*98*), an open-source, in-browser analysis program with built-in tools including SegmentAnything and Piscis. In NimbusImage, cell cytoplasm (“cell annotations”) was manually segmented using the “Manual Blob Annotation” tool, and cell nuclei (“nucleus annotations”) were segmented using the SegmentAnything Model (https://github.com/facebookresearch/segment-anything). Morphology measurements for these annotations were calculated using the “Blob Metrics” tool. *Pycard* RNA-FISH spots were identified across all Z-planes using the deep learning algorithm, Piscis (*99*), and linked to cell annotations using the “Point Count 3D Projection” tool.

Fluorescence intensities for the *Pycard* RNA-FISH and RPS6 IF signals were calculated using the “Blob Intensity Measurements” tool. Within each image frame, a region without cells was also annotated to assess background fluorescence intensity. For cell annotations, the mean fluorescence intensity across the annotation was calculated across all 5 Z-planes, and the maximum value was selected. These values were then background corrected within the image frame. For nucleus annotations, the mean perinuclear fluorescence intensity was calculated in a 10-pixel ring around the annotation (excluding the annotation area itself) across all 5 Z-planes, and the maximum value was selected. These values were then background corrected within the image frame.

*Pycard* RNA-FISH spots per cell annotation were calculated using the “Point Count 3D Projection” tool and normalized by cell annotation area (*100*). To count perinuclear *Pycard* RNA-FISH spots, we used the JSON outputs of NimbusImage nucleus and Piscis annotations and the *sf* package in R to identify all spots located within a 10-pixel buffer of the nucleus annotation across all 5 Z-planes.

Data and code are accessible at: https://upenn.box.com/s/aklb7gogtzl5kzjl0vjijjtlxruyjc3x.

### Statistical Analysis

Unless otherwise noted, all data points are shown as mean values of biological replicates and were pooled from at least 3 independent experiments, with error bars representing the SD. The sample size and number of independent experiments are indicated in each figure legend. All statistical tests are described in the figure captions and were performed using Prism (GraphPad). *P* values of 0.05 or less were considered statistically significant.

## SUPPLEMENTAL MATERIALS AND METHODS

### Metabolomics

After 20-24 hours of culture in complete or BCAA-free medium, BMDMs (1 x 10^6^ per sample) were washed with PBS and flash frozen using −80°C methanol. Cell pellets were extracted in a 1 mL, −80°C solution containing 80% ethanol, 20% Optima water, and a mix of 13C/15N-labeled amino acid internal standards (Cambridge Isotope Laboratories #MSKA2-1.2). Samples were sonicated on ice using a sonic dismembrator (ThermoFisher Scientific) 30 times over 15 seconds and then incubated on ice for 10 minutes. Debris was pelleted at 12,000 x g for 10 minutes at 4°C, and the solvent was dried under nitrogen gas using a blowdown evaporator (Organomation #11634). Dry metabolites were re-suspended in 300 μL of a 5% methanol solution, vortexed for 1 minute, dissolved using an ultrasonic bath for 15 minutes, spun down at 12,000 x g for 10 minutes at 4°C, and then distributed to HPLC vials for LC-HRMS analysis. A pooled QC sample was generated for monitoring intra-run variance by mixing 20 μL of each resuspended experimental sample within the run.

Metabolites were separated with a 150 mm x 2.1 mm (1.9 μm particle size) Hypersil GOLD HPLC column (ThermoFisher Scientific #25402-152130) using an UltiMate 3000 UHPLC (ThermoFisher Scientific) equipped with a refrigerated autosampler (6°C) and column heater (55°C). The solvent was methanol with 0.1% formic acid, and the gradient was arranged as follows: 0.5% solvent at 0 minutes, 0.5% solvent at 2 minutes, 50% solvent at 6 minutes, 100% solvent at 12 minutes, 100% solvent at 16 minutes, and back to 0.5% solvent at 17 minutes, maintained for 3 more minutes for column re-equilibration. The flow rate was 0.45 mL/minute. A Q Exactive HF mass analyzer (ThermoFisher Scientific) equipped with a heated electrospray ionization source was operated in positive mode in full scan at 120,000 resolution, automatic gain control (AGC) target=1 x 10^6^, maximum ion injection time (IT)=100 milliseconds, and scan range=70 to 800 m/z. The pooled QC samples were used for metabolite identification by generating MS/MS spectra (ddMS2) of the top 10 features at 15,000 resolution, AGC target=1 x 10^5^, maximum IT=25 milliseconds, and (N)CE/stepped NCE=30, 50, 60 v.

For data analysis, metabolites were identified based on the exact mass +/- 5 ppm and retention time of authentic standards (Cambridge Isotope Laboratories #MSK-A2-1.2). Peak integration was calculated using Xcalibur 4.2 (ThermoFisher Scientific). Normalized metabolite abundances were generated by plotting the ratio of the detected metabolite peak area relative to its corresponding 13C-labeled internal standard, or, if not present, the internal standard with the nearest retention time.

### RNA and Protein Stability Assays

BMDMs were seeded in 24-well, tissue culture-treated plates at 2.5 x 10^5^ cells/mL (500 µL per well). After 20-24 hours of culture in complete or BCAA-free medium, cells were stimulated with 100 ng/mL LPS for 2 hours. Medium was then aspirated and replaced with fresh complete or BCAA-free medium containing 10 µg/mL actinomycin D or cycloheximide to inhibit RNA transcription or protein translation, respectively, along with the pan-caspase inhibitor Z-VAD-FMK (30 µM) to limit cell death associated with these treatments. Cell lysates were collected at the indicated timepoints for RT-qPCR and Western blot analyses of mRNA and protein decay.

### Puromycin Labeling of Newly Synthesized Proteins

BMDMs were seeded onto 6-well tissue culture-treated plastic plates at a concentration of 2.5 x 10^5^ cells/mL in a volume of 4 mL. After 20-24 hours of culture in complete or BCAA-free medium, BMDMs were treated with 100 ng/mL LPS or vehicle for 3.5 hours prior to pulsing with 10 µg/mL puromycin (Cayman Chemical #13884). Cell lysates were harvested after 30 minutes, and samples were prepared for Western blot.

## ACKNOWLEDGMENTS

The authors thank Drs. Yangzhu Du, Honghong Sun, and Eline Luning Prak of the Human Immunology Core of the Penn Center for AIDS Research and Abramson Cancer Center at the University of Pennsylvania Perelman School of Medicine for assistance with Luminex assay. We also thank the Human Immunology Core for providing purified primary human monocytes. The Human Immunology Core is supported in part by National Institutes of Health grants P30AI045008 and P30CA016520 (Human Immunology Core RRID: SCR_022380). We thank Gordon Ruthel of the Penn Vet Imaging Core for assistance with microscopy. Lastly, we thank all members of the Sunny Shin, Will Bailis, and Igor Brodsky Laboratories at the University of Pennsylvania and Children’s Hospital of Philadelphia for providing feedback and support. Schematics were generated using BioRender.

## Supplemental Figures

**Fig. S1:** Cytotoxicity was determined by quantifying LDH release after culturing BMDMs for 20-24 hours in (**A**) 100% amino acids (Control) or 10% amino acids (10% AAs) media or (**D**) custom-formulated media containing all amino acids at standard concentrations (Control) or lacking a single amino acid (Table S1), as indicated. Release of (**B**) LDH and (**C**) IL-1β was quantified by cytotoxicity assay and ELISA, respectively, after LPS priming (100 ng/mL, 4 hours) and subsequent nigericin treatment (5 µM, 1 hour), either immediately following (Immediate Restriction, IR) or after 20-24 hours of culture (Prolonged Restriction, PR) in control or 10% AAs media. (**E**) Intracellular leucine, isoleucine, and valine levels were quantified by mass spectrometry of BMDM lysates after 20-24 hours of culture in a BCAA-free RPMI 1640 medium supplemented with or without leucine, isoleucine, and valine at standard concentrations. Release of (**F**) LDH and (**G**) IL-1β was quantified by cytotoxicity assay and ELISA, respectively, after LPS priming (100 ng/mL, 4 hours) and subsequent vehicle or nigericin (5-10 µM) treatment (50-120 minutes) immediately following BMDM culture in BCAA-free RPMI 1640 medium supplemented with or without leucine, isoleucine, and valine at standard concentrations. Release of (**H**) LDH and (**I**) IL-1β was quantified from BMDMs primed with LPS (100 ng/mL, 4 hours) and treated with nigericin (5 µM, 45-150 minutes) after 24 hours of culture in BCAA-free RPMI 1640 medium supplemented with or without leucine, isoleucine, and valine at standard concentrations, as indicated by “+” or “-” at “T=-24 h”. “T=0 h” indicates the culture medium (“+” or “-” BCAAs) added to BMDMs at the time of LPS priming. (**J**) TNF and (**K**) IL-6 release was quantified by ELISA of supernatants collected from LPS-treated (100 ng/mL, 4 hours) BMDMs after immediate or prolonged (i.e., 20-24 hours) culture in BCAA-containing or BCAA-free RPMI 1640 medium. (**L**) Cleavage of caspase-1 and GSDMD was assessed by Western blot of lysates after LPS priming (100 ng/mL, 4 hours) and subsequent vehicle or nigericin (10 µM) treatment (45 minutes) of BMDMs immediately following culture in BCAA-free RPMI 1640 medium supplemented with or without leucine, isoleucine, and valine at standard concentrations. Data represent (A to D and F to K) the mean values of biological duplicates or triplicates pooled from ≥3 independent experiments +/- SD, (E) the average metabolite concentration quantified from 5 replicate wells of BMDMs per experiment across 2 independent experiments +/- SD, and (L) results from 1 experiment representative of 3 independent experiments. Statistical significance was determined using (B, C, H and I) one-way ANOVA and (F, G, J, and K) two-way ANOVA. FL=full length. ns=not significant. ND=not detectable. *p < 0.05 **p < 0.01 ***p < 0.001 ****p < 0.0001

**Fig. S2:** hMDMs from 3-4 independent donors were cultured for (**A**) and (**B**) 24 hours or (**C**) 24-48 hours in BCAA-free RPMI 1640 medium supplemented with or without leucine, isoleucine, and valine at standard concentrations before priming with LPS (100 ng/mL) or vehicle control for 4 hours. (**A**) LDH and (**B**) human IL-1β (hIL-1β) release was quantified by cytotoxicity assay and ELISA, respectively, after subsequent treatment of hMDMs with vehicle control or nigericin (60 µM) for 3-4 hours. (**C**) Lysates were collected for Western blot analysis of pro-IL-1β protein. (**D**) and (**E**) BMDMs were cultured for 20-24 hours in BCAA-free RPMI 1640 medium supplemented with or without leucine, isoleucine, and valine at standard concentrations before infecting with *Legionella pneumophila* (LP02 Δ*flaA*, MOI=50). (**D**) LDH release was measured by cytotoxicity assay at 8 hours post-infection, and (**E**) IL-1β release was measured by ELISA at 8 and 18 hours post-infection. (**F**) to (**H**) BMDMs were cultured for 20-24 hours in BCAA-free RPMI 1640 medium supplemented with or without leucine, isoleucine, and valine at standard concentrations before priming with LPS (100 ng/mL) for 6 hours and then cytosolically delivering LPS (1 µg/mL) via transfection with FuGENE HD for 5 hours. Supernatants were collected for quantification of (**F**) LDH release by cytotoxicity assay as well as (**G**) IL-1β and (**H**) IL-1α release by ELISA. (**I**) Schematic of LPS endotoxic shock model after dietary BCAA restriction *in vivo*. (**J**) 6-7-week-old mice were placed on a complete (Control) amino acid diet or BCAA-restricted (25% BCAAs) diet for 5 weeks before initiating endotoxemia. Peritoneal wash was collected 3 hours after 10 mg/kg intraperitoneal (i.p.) LPS injection, and cytokine/chemokine levels were measured by Luminex assay. Data represent (A, B, and D to H) the mean values of biological duplicates or triplicates pooled from ≥3 independent experiments +/- SD, (C) Western blot results of hMDMs derived from 4 independent donors, and (J) cytokine or chemokine values +/- SD, where each data point represents the value quantified from an individual mouse (n=10 males on a complete diet, n=9 males on a BCAA-restricted diet). Statistical significance was determined using (A, E, G, and H) two-way ANOVA, (B, D, and F) unpaired student’s t-test, and (J) Mann-Whitney test. The dotted line in (C) represents splicing together bands from 2 non-adjacent lanes run on the same gel. ns=not significant. ND=not detectable. OOR=out of range. *p < 0.05 **p < 0.01 ***p < 0.001 ****p < 0.0001

**Fig. S3:** (**A**) to (**C**) BMDMs were cultured in BCAA-free RPMI 1640 medium supplemented with or without leucine, isoleucine, and valine at standard concentrations and then immediately primed with LPS (100 ng/mL) or vehicle control for 4 hours. Lysates were collected for RT-qPCR analysis of (**A**) *Il1b* and (**B**) *Nlrp3* mRNA or (**C**) Western blot analysis of pro-IL-1β, NLRP3, ASC, and caspase-1 protein. (**D**) to (**H**) BMDMs were cultured in BCAA-free RPMI 1640 medium supplemented with or without leucine, isoleucine, and valine at standard concentrations for 20-24 hours and then treated with LPS (100 ng/mL). After 2 hours (h) of LPS priming, BMDMs were either pulsed with (**D**) to (**G**) 10 µg/mL actinomycin D (Act D) or (**H**) 10 µg/mL cycloheximide (CHX) to halt RNA transcription and protein translation, respectively. Lysates were then collected at the indicated timepoints for RT-qPCR analysis of (**D**) *Il1b*, (**E**) *Nlrp3*, (**F**) *Pycard*, and (**G**) *Casp1* mRNA or (**H**) Western blot analysis of pro-IL-1β, NLRP3, ASC, and caspase-1 protein. Data represent (A, B, and D to G) the mean values of biological duplicates pooled from ≥3 independent experiments +/- SD and (C and H) results from 1 experiment representative of 3 independent experiments. Statistical significance was determined using (A and B) two-way ANOVA and (D to G) unpaired student’s t-test. ns=not significant. *p < 0.05 **p < 0.01 ***p < 0.001 ****p < 0.0001

**Fig. S4:** (**A**) to (**J**) BMDMs were cultured in BCAA-free RPMI 1640 medium supplemented with or without leucine, isoleucine, and valine at standard concentrations for 20-24 hours. (**A**) After 4 hours of priming with LPS (100 ng/mL) or vehicle control, lysates were collected for polysome profiling with the abundance of monosome-associated RNA calculated by area under the curve (AUC) analysis. (**B**) After 3.5 hours of priming with LPS (100 ng/mL) or vehicle control, BMDMs were pulsed with 10 µg/mL puromycin for 30 minutes, and lysates were collected for Western blot analysis of puromycin incorporation. (**C**) to (**H**) After 4 hours of priming with LPS (100 ng/mL) or vehicle control, lysates were collected for polysome profiling. (**C**) *Il1a* mRNA levels were quantified in unfractionated whole-cell lysate (Input) samples and (**D**) in 60S, monosome, disome, and polysome fractions, with expression normalized to an equivalent amount of *Luciferase* mRNA spiked into each sample. (**E**) Polysome-associated *Il1a* levels were measured by AUC analysis of fractions 8 to 13 (F8-F13 in [D]). (**F**) *Casp11* mRNA levels were quantified in unfractionated whole-cell lysate (Input) samples and (**G**) in 60S, monosome, disome, and polysome fractions, with expression normalized to *Luciferase* mRNA. (**H**) Polysome-associated *Casp11* levels were measured by AUC analysis of fractions 8 to 13 (F8-F13 in [G]). (**I**) After 6 hours of LPS (100 ng/mL) priming, BMDM lysates were collected for Western blot analysis of caspase-11 protein, with (**J**) optical density (OD) analysis of caspase-11 expression normalized to vehicle-treated control (Ctrl.) BMDMs that were cultured in complete medium (CM). Data represent (A, E, and H) AUC data pooled from 3 independent experiments +/- SD, (B) results from 3 independent experiments, (C and F) the mean values of technical triplicates pooled from 3 independent experiments +/- SD, (D and G) RT-qPCR results showing technical triplicates +/- SD from 1 experiment representative of 3 independent experiments, (I) results from 1 experiment representative of 4 independent experiments, and (J) normalized densitometry values pooled from 4 independent experiments +/- SD. (A, C, E, F, H, and J) Statistical significance was determined using two-way ANOVA. ns=not significant. *p < 0.05 **p < 0.01

**Fig. S5:** (**A**) Schematic representation of the catabolic and signaling properties of the BCAAs. (**B**) to (**M**) BMDMs were cultured for 20-24 hours in BCAA-free RPMI 1640 medium supplemented with or without leucine, isoleucine, and valine at standard concentrations before priming with LPS (100 ng/mL) or vehicle control for 4 hours. (**B**) DBT wild-type (WT) and knockout (KO) BMDM lysates were collected for Western blot analysis of DBT and pro-IL-1β protein. Release of (**C**) IL-1β and (**D**) LDH was quantified from LPS-primed DBT WT and KO BMDMs via ELISA and cytotoxicity assay, respectively, after subsequent treatment with nigericin (5 µM) for 40 minutes. (**E**) WT BMDM lysates were collected for Western blot analysis of ATF4. (**F**) to (**H**) WT BMDM lysates were collected for polysome profiling. (**F**) *Atf4* mRNA levels were quantified in unfractionated whole-cell lysate (Input) samples and (**G**) in 60S, monosome, disome, and polysome fractions, with expression normalized to an equivalent amount of *Luciferase* mRNA spiked into each sample. (**H**) Polysome-associated *Atf4* levels were measured by area under the curve (AUC) analysis of fractions 8 to 13 (F8-F13 in [G]) and normalized to vehicle-treated control (Ctrl.) BMDMs that were cultured in complete medium (CM). (**I**) and (**J**) GCN2iB (1 µM) or vehicle control was added to WT BMDMs for 20-24 hours parallel to BCAA starvation. (**I**) Lysates were collected for Western blot analysis of ATF4 and pro-IL-1β protein, and (**J**) LDH release into supernatants was quantified via cytotoxicity assay after subsequent treatment of LPS-primed BMDMs with nigericin (5 µM) for 45-60 minutes. (**K**) GCN2 WT and KO BMDM lysates were collected for Western blot analysis of GCN2, ATF4, and pro-IL-1β protein. Release of (**L**) LDH and (**M**) IL-1β was quantified from LPS-primed GCN2 WT and KO BMDMs via cytotoxicity assay and ELISA, respectively, after subsequent treatment with nigericin (5 µM) for 40-95 minutes. Data represent (B) results from 1 experiment representative of 2 independent experiments, (C and D) biological triplicates +/- SD from 1 experiment representative of 2 independent experiments, (E and I) results from 1 experiment representative of 3 independent experiments, (F) the mean values of technical triplicates pooled from 3 independent experiments +/- SD, (G) RT-qPCR results showing technical triplicates +/- SD from 1 experiment representative of 3 independent experiments, (H) normalized AUC data pooled from 3 independent experiments +/- SD, (J, L, and M) the mean values of biological duplicates or triplicates pooled from ≥3 independent experiments +/- SD, and (K) results from 1 experiment representative of 2-3 independent experiments. (L) Zero and 100% LDH release (from untreated and detergent-permeabilized cells, respectively) were calculated separately for GCN2 WT and KO BMDMs. Statistical significance was determined using (C, D, F, H, and J) two-way ANOVA and (L and M) unpaired student’s t-test. ns=not significant. *p < 0.05 **p < 0.01 ***p < 0.001 ****p < 0.0001

**Fig. S6:** (**A**) to (**C**) BMDMs were pre-treated with Torin 1 (1 µM) or left untreated (Control) for 30 minutes prior to LPS priming (100 ng/mL, 4 hours), and lysates were collected for polysome profiling. (**A**) Total RNA absorbance was quantified across sucrose gradient fractions. (**B**) and (**C**) RNA sequencing was performed on unfractionated whole-cell lysates (Input) or pooled polysome fractions, and upregulated/downregulated differentially expressed genes (DEGs) were analyzed in and compared across each sample type. (**D**) to (**F**) BMDMs were cultured for 20-24 hours in BCAA-free RPMI 1640 medium supplemented with or without leucine, isoleucine, and valine at standard concentrations before priming with LPS (100 ng/mL) or vehicle control for 4 hours. (**D**) *Casp1* mRNA levels were quantified in unfractionated whole-cell lysate (Input) samples and (**E**) in 60S, monosome, disome, and polysome fractions, with expression normalized to an equivalent amount of *Luciferase* mRNA spiked into each sample. (**F**) Polysome-associated *Casp1* levels were measured by area under the curve (AUC) analysis of fractions 8 to 13 (F8-F13 in [E]) and normalized to vehicle-treated control (Ctrl.) BMDMs that were cultured in complete medium (CM). Data represent (A) results from 1 experiment representative of 3 independent experiments, (B and C) values averaged from analysis of samples collected across 3 independent experiments, (D) the mean values of technical triplicates pooled from 3 independent experiments +/- SD, (E) RT-qPCR results showing technical triplicates +/- SD from 1 experiment representative of 3 independent experiments, and (F) normalized AUC data pooled from 3 independent experiments +/- SD. (D and F) Statistical significance was determined using two-way ANOVA. ns=not significant.

**Fig. S7:** (**A**) to (**E**) BMDMs were cultured for 20-24 hours in BCAA-free RPMI 1640 medium supplemented with or without leucine, isoleucine, and valine at standard concentrations before priming with LPS (100 ng/mL) or vehicle control for 4 hours. (**A**) to (**D**) Untreated, LPS-primed, and LPS-primed plus nigericin-stimulated (5 µM, 20-30 minutes) ASC-citrine BMDMs were fixed for immunofluorescence and RNA-FISH labeling of RPS6 and *Pycard* mRNA, respectively. Quantification of maximum cellular (**A**) RPS6 and (**B**) *Pycard* mRNA fluorescence intensity, (**C**) *Pycard* mRNA spot counts within a 10-pixel buffer of the nucleus annotation (Perinuclear Space), and (**D**) cellular *Pycard* mRNA spot counts, normalized to cell area. (**E**) For the final 20, 40, or 60 minutes of LPS priming, cycloheximide (CHX, 50 µg/mL) was added to halt protein translation in ASC-citrine BMDMs, and lysates were collected for Western blot analysis of NLRP3, ASC, and caspase-1 protein. (**F**) Schematic of the proposed model in which environmental BCAA sensing licenses mTOR-driven *Il1b* transcription and ASC translation, which accelerates NLRP3 inflammasome assembly. Data represent (A to D) violin plots showing the median, interquartile range, and full range of the data collected from 2 independent experiments (>545 cells/nuclei analyzed in total for each condition) and (E) results from 1 experiment representative of 3 independent experiments. (A to D) Statistical significance was determined using two-way ANOVA. ns=not significant. *p < 0.05 **p < 0.01 ***p < 0.001 ****p < 0.0001

