## Supplemental Figures and Table for "Environmental Amino Acid Sensing Regulates the Rate of ASC Translation and NLRP3 Inflammasome Assembly"

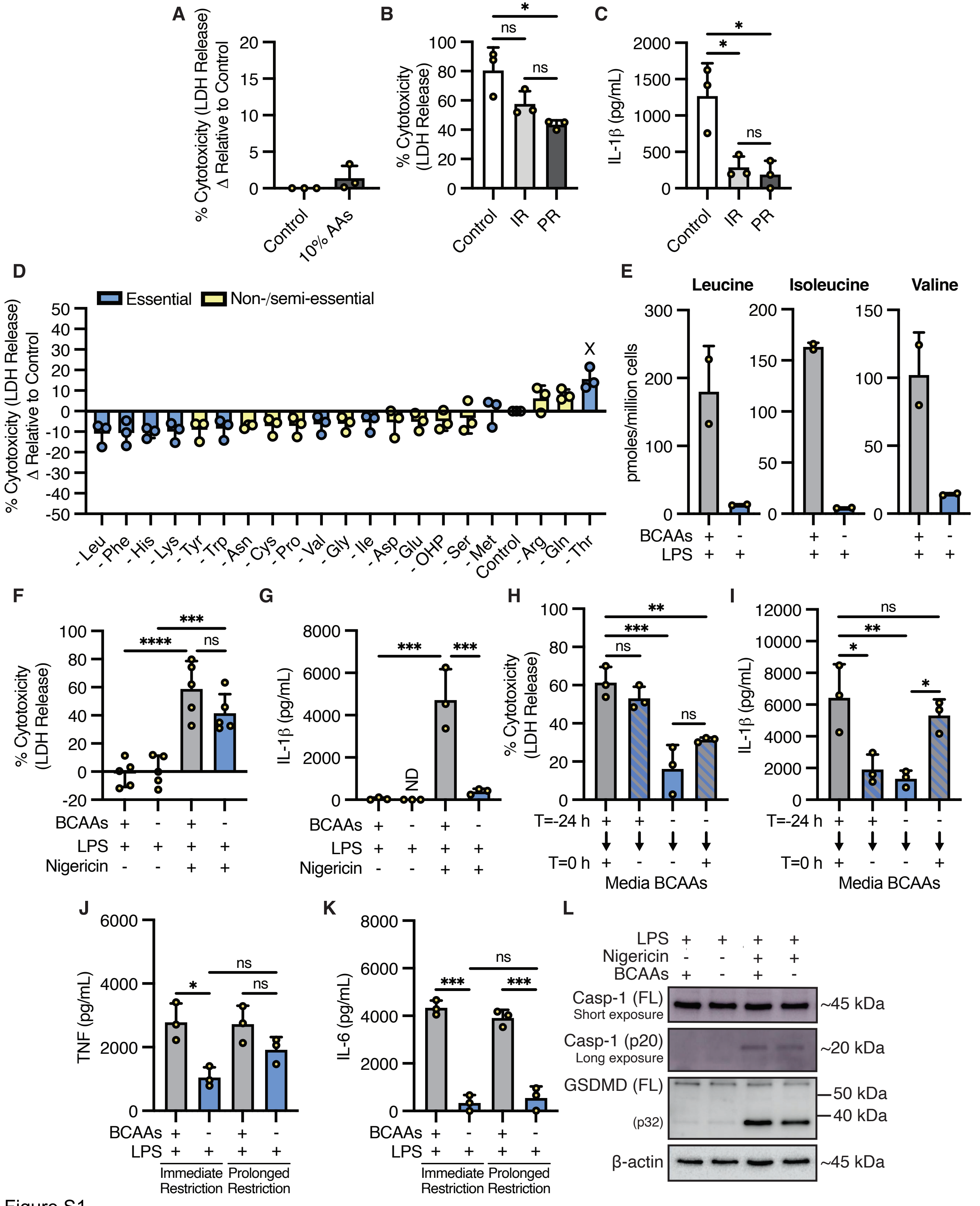
A
% Cytotoxicity (LDH Release) Δ Relative to Control
Control
10% AAs
B
% Cytotoxicity (LDH Release)
Control
IR
PR
ns
*
C
IL-1β (pg/mL)
D
Essential
Non-/semi-essential
X
-Leu -Phe -His -Lys -Tyr -Trp -Asn -Cys -Pro -Val -Gly -Ile -Asp -Glu -OHP -Ser -Met Control -Arg -Gln -Thr
E
Leucine
Isoleucine
Valine
pmoles/million cells
BCAAs
LPS
F
G
ND
Nigericin
H
I
T=-24 h
T=0 h
Media BCAAs
J
TNF (pg/mL)
Immediate Restriction
Prolonged Restriction
K
IL-6 (pg/mL)
L
Casp-1 (FL)
Short exposure
Casp-1 (p20)
Long exposure
GSDMD (FL)
(p32)
β-actin
~45 kDa
~20 kDa
50 kDa
40 kDa
Figure S1

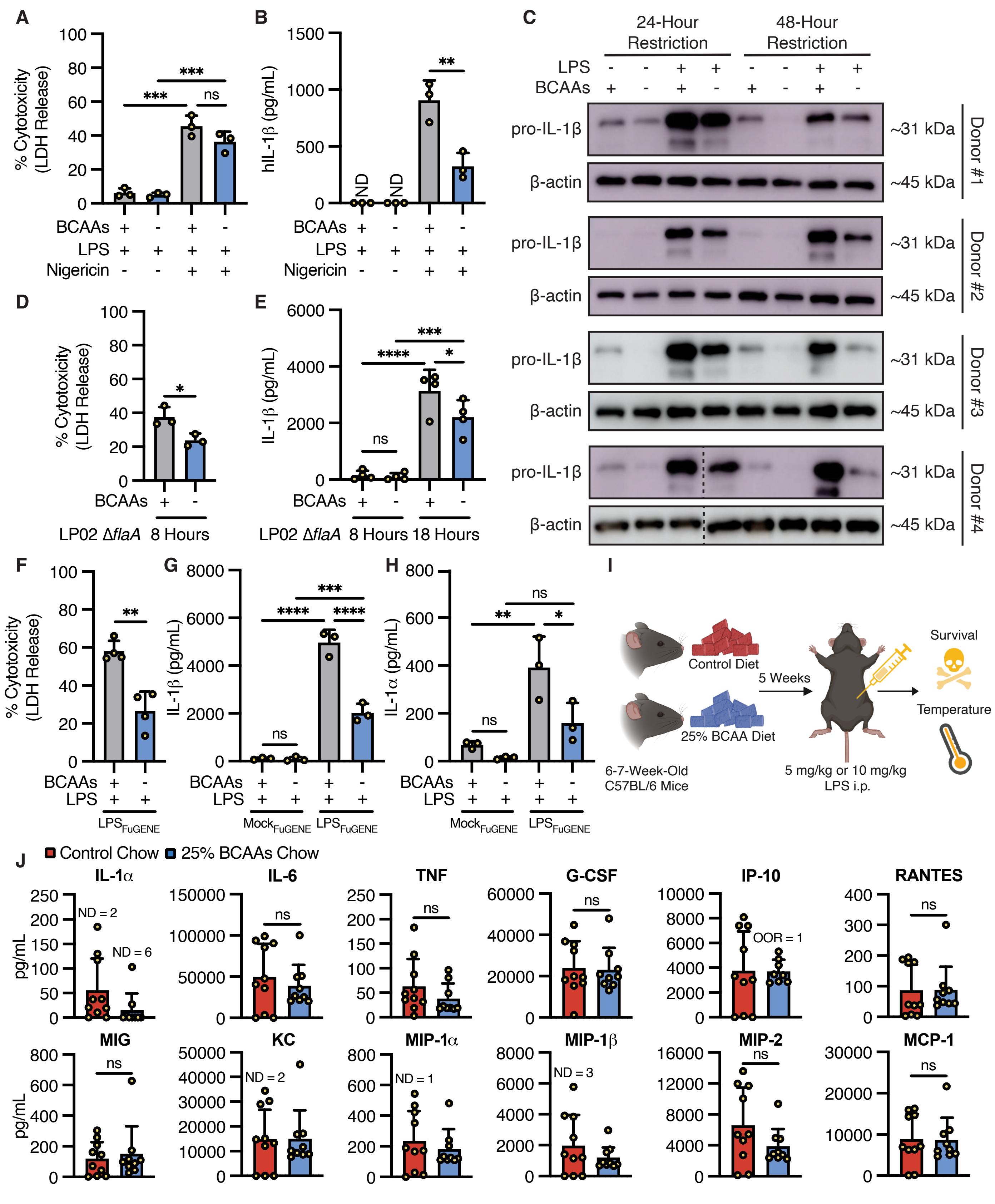
A
% Cytotoxicity (LDH Release)
BCAAs
LPS
Nigericin
B
hIL-1β (pg/mL)
ND
C
24-Hour Restriction
48-Hour Restriction
pro-IL-1β
β-actin
~31 kDa
~45 kDa
Donor #1
Donor #2
Donor #3
Donor #4
D
E
IL-1β (pg/mL)
LP02 ΔflaA
8 Hours
18 Hours
F
G
H
IL-1α (pg/mL)
LPS_FuGENE
Mock_FuGENE
I
Control Diet
25% BCAA Diet
5 Weeks
Survival
Temperature
6-7-Week-Old C57BL/6 Mice
5 mg/kg or 10 mg/kg LPS i.p.
J
Control Chow
25% BCAAs Chow
IL-1α
IL-6
TNF
G-CSF
IP-10
RANTES
MIG
KC
MIP-1α
MIP-1β
MIP-2
MCP-1
pg/mL
ND = 2
ND = 6
OOR = 1
ND = 2
ND = 1
ND = 3
ns

Figure S2

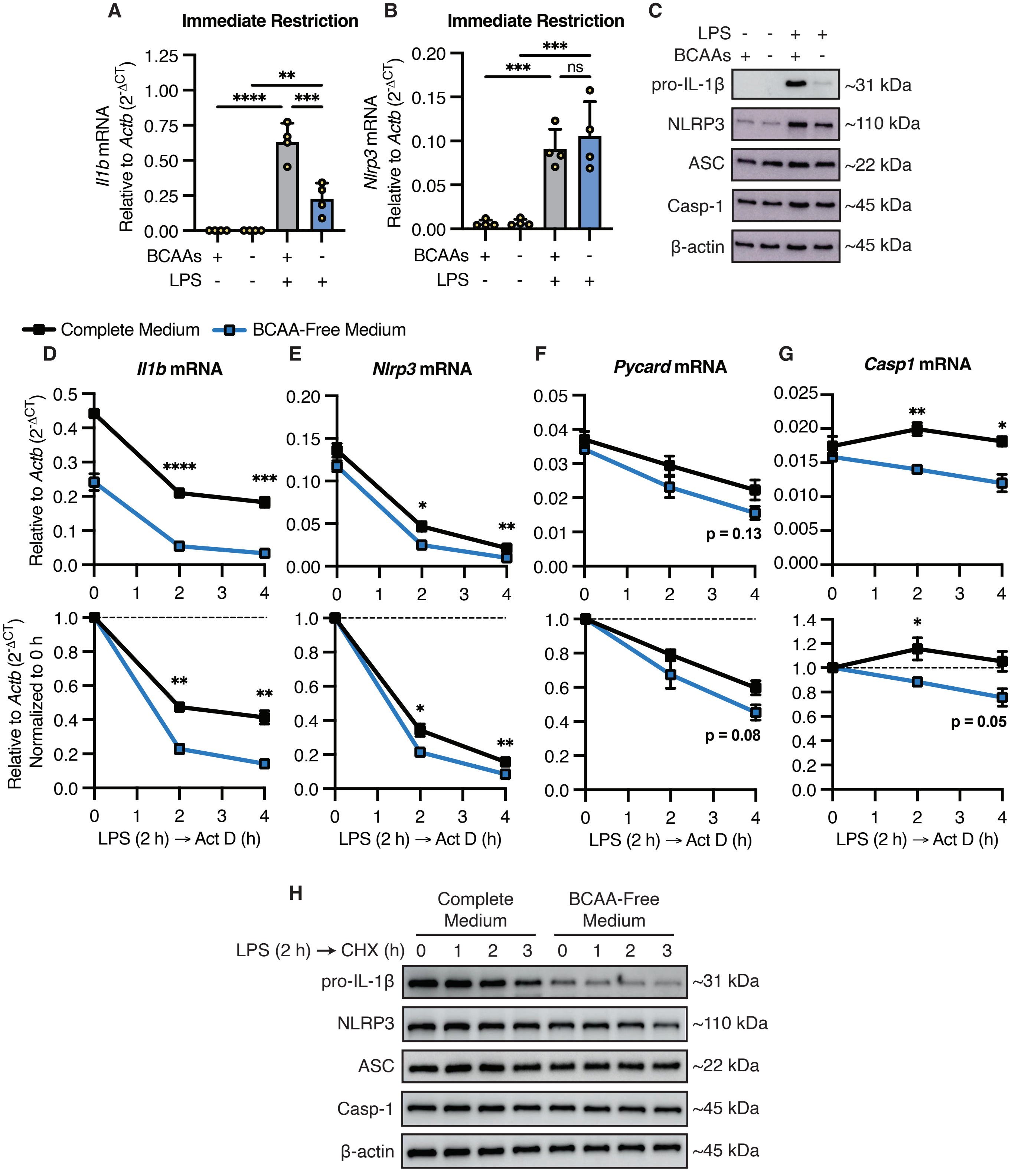

A
Immediate Restriction
Il1b mRNA Relative to Actb (2-ΔCT)
BCAAs + - + -
LPS - - + +
B
Immediate Restriction
Nlrp3 mRNA Relative to Actb (2-ΔCT)
ns
C
LPS - - + +
BCAAs + - + -
pro-IL-1β ~31 kDa
NLRP3 ~110 kDa
ASC ~22 kDa
Casp-1 ~45 kDa
β-actin ~45 kDa
Complete Medium
BCAA-Free Medium
D
Il1b mRNA
E
Nlrp3 mRNA
F
Pycard mRNA
p = 0.13
p = 0.08
G
Casp1 mRNA
p = 0.05
Relative to Actb (2-ΔCT)
Relative to Actb (2-ΔCT) Normalized to 0 h
LPS (2 h) → Act D (h)
H
Complete Medium
BCAA-Free Medium
LPS (2 h) → CHX (h) 0 1 2 3 0 1 2 3
pro-IL-1β ~31 kDa
NLRP3 ~110 kDa
ASC ~22 kDa
Casp-1 ~45 kDa
β-actin ~45 kDa


Figure S3

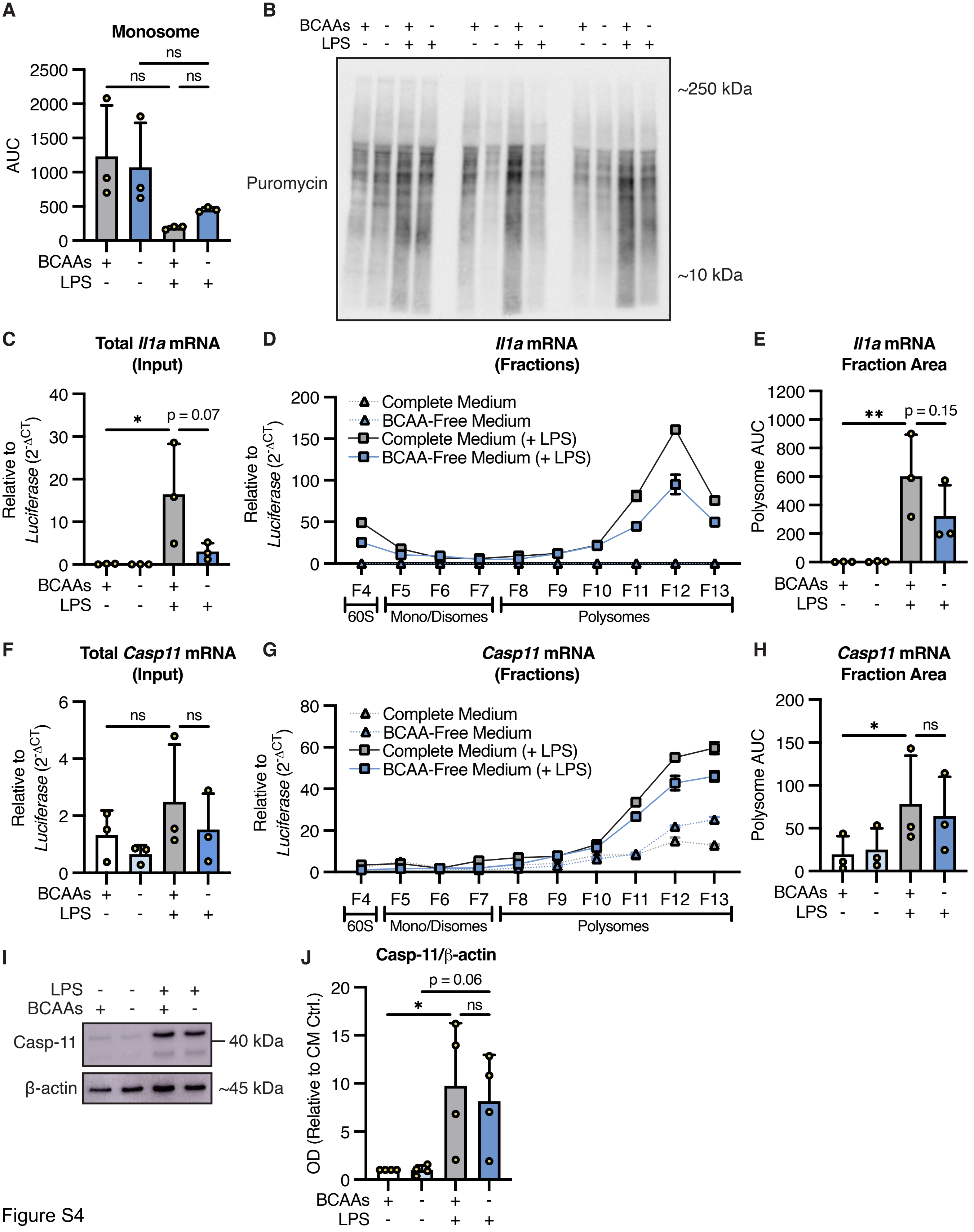

A
Monosome
AUC
ns
BCAAs
LPS
B
Puromycin
~250 kDa
~10 kDa
C
Total Il1a mRNA (Input)
Relative to Luciferase (2^-ΔCT)
*
p = 0.07
D
Il1a mRNA (Fractions)
Complete Medium
BCAA-Free Medium
Complete Medium (+ LPS)
BCAA-Free Medium (+ LPS)
F4 F5 F6 F7 F8 F9 F10 F11 F12 F13
60S
Mono/Disomes
Polysomes
E
Il1a mRNA Fraction Area
Polysome AUC
**
p = 0.15
F
Total Casp11 mRNA (Input)
G
Casp11 mRNA (Fractions)
H
Casp11 mRNA Fraction Area
I
Casp-11
β-actin
40 kDa
~45 kDa
J
Casp-11/β-actin
OD (Relative to CM Ctrl.)
p = 0.06


Figure S4

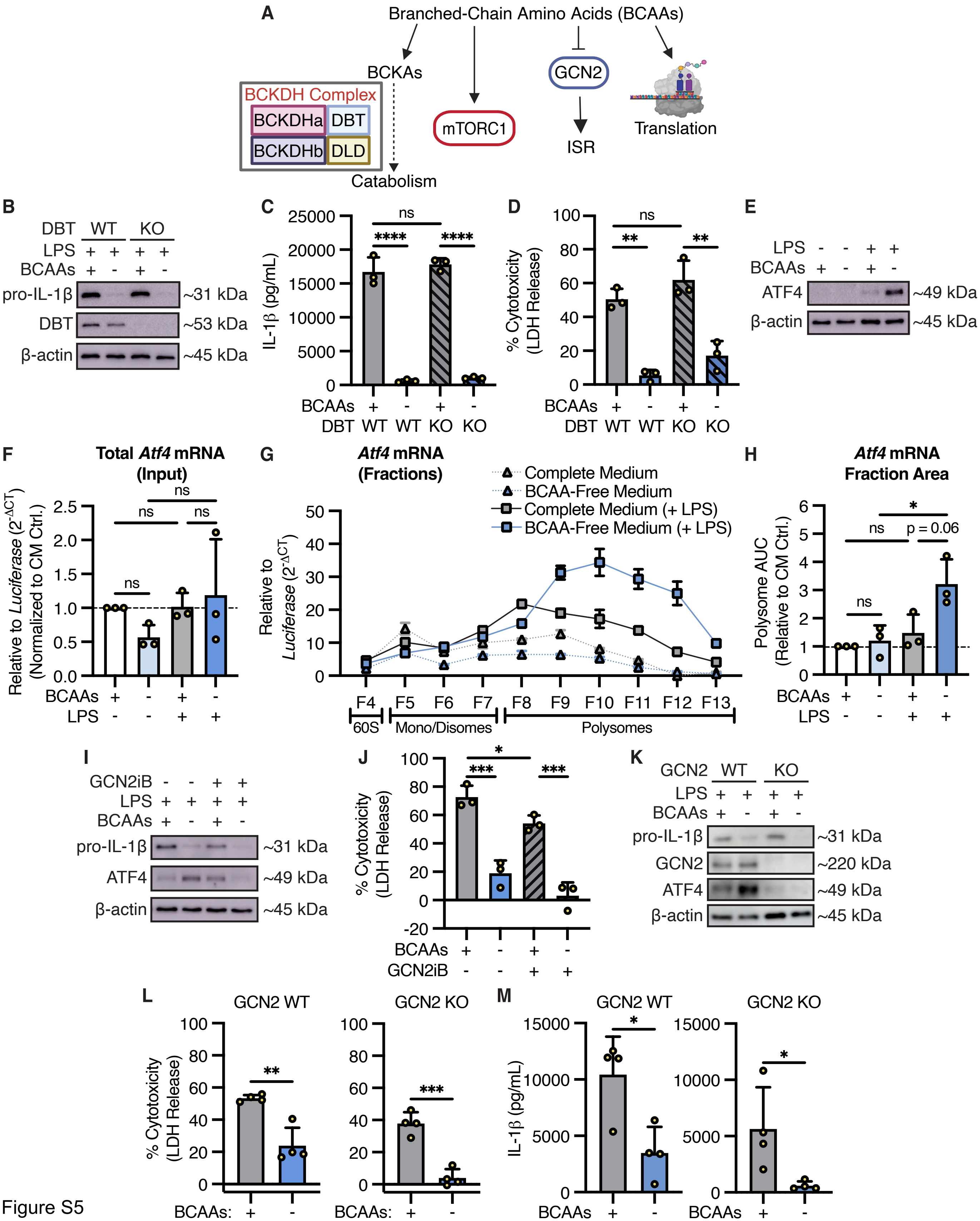

A
Branched-Chain Amino Acids (BCAAs)
BCKAs
BCKDH Complex
BCKDHa
DBT
BCKDHb
DLD
Catabolism
mTORC1
GCN2
ISR
Translation
B
DBT WT KO
LPS + + + +
BCAAs + - + -
pro-IL-1β ~31 kDa
DBT ~53 kDa
β-actin ~45 kDa
C
IL-1β (pg/mL)
BCAAs + - + -
DBT WT WT KO KO
D
% Cytotoxicity (LDH Release)
E
LPS - - + +
BCAAs + - + -
ATF4 ~49 kDa
β-actin ~45 kDa
F
Total Atf4 mRNA (Input)
Relative to Luciferase (2^-ΔCT) (Normalized to CM Ctrl.)
G
Atf4 mRNA (Fractions)
Complete Medium
BCAA-Free Medium
Complete Medium (+ LPS)
BCAA-Free Medium (+ LPS)
Relative to Luciferase (2^-ΔCT)
F4 F5 F6 F7 F8 F9 F10 F11 F12 F13
60S
Mono/Disomes
Polysomes
H
Atf4 mRNA Fraction Area
Polysome AUC (Relative to CM Ctrl.)
p = 0.06
I
GCN2iB - - + +
LPS + + + +
BCAAs + - + -
pro-IL-1β ~31 kDa
ATF4 ~49 kDa
β-actin ~45 kDa
J
% Cytotoxicity (LDH Release)
BCAAs + - + -
GCN2iB - - + +
K
GCN2 WT KO
LPS + + + +
BCAAs + - + -
pro-IL-1β ~31 kDa
GCN2 ~220 kDa
ATF4 ~49 kDa
β-actin ~45 kDa
L
GCN2 WT
GCN2 KO
% Cytotoxicity (LDH Release)
BCAAs: + -
M
GCN2 WT
GCN2 KO
IL-1β (pg/mL)
BCAAs + -


Figure S5

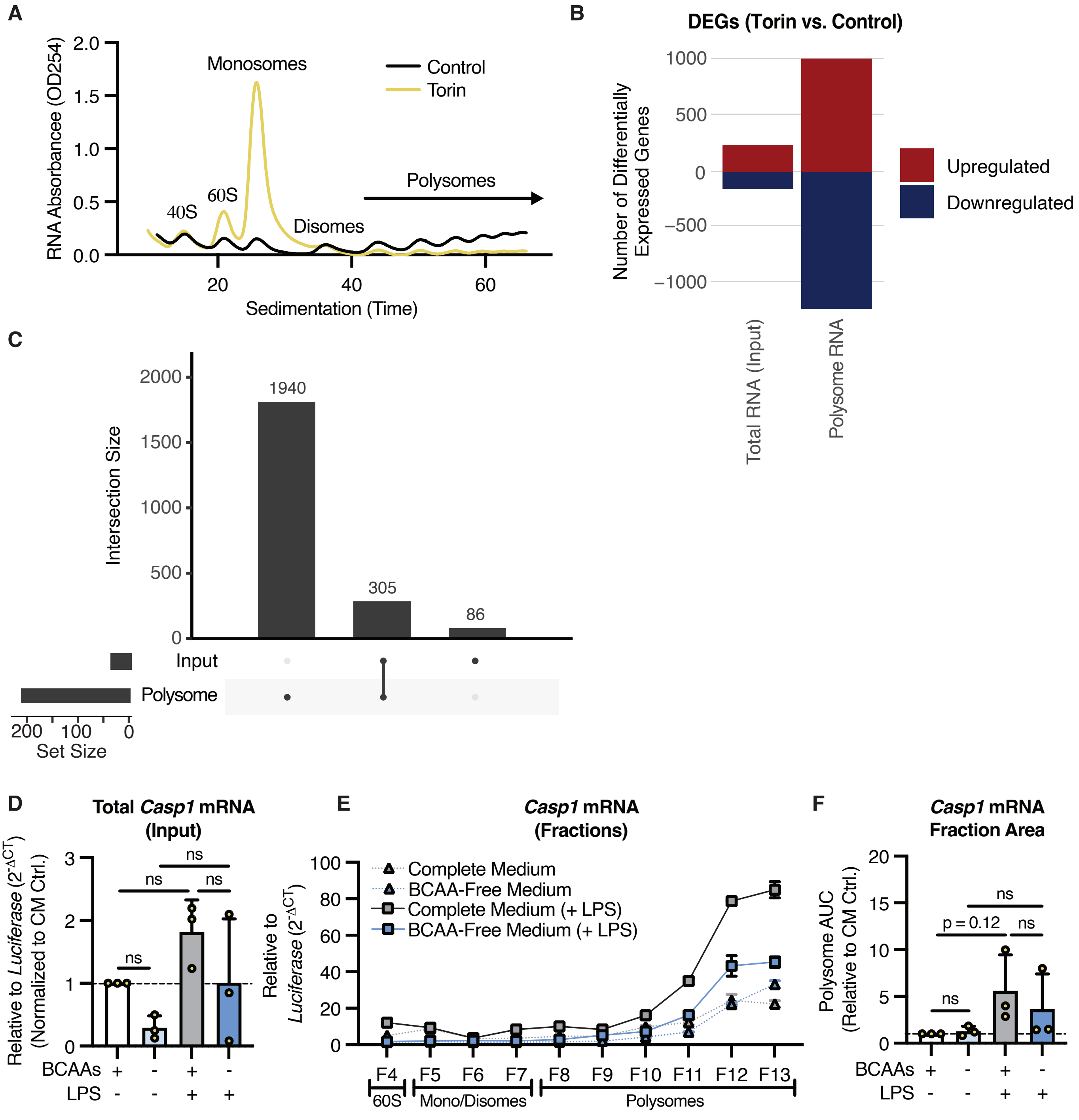
A
RNA Absorbancee (OD254)
2.0
1.5
1.0
0.5
0.0
Monosomes
Control
Torin
40S
60S
Disomes
Polysomes
20
40
60
Sedimentation (Time)
B
DEGs (Torin vs. Control)
Number of Differentially Expressed Genes
1000
500
0
−500
−1000
Upregulated
Downregulated
Total RNA (Input)
Polysome RNA
C
Intersection Size
2000
1500
1000
500
0
1940
305
86
Input
Polysome
200
100
0
Set Size
D
Total Casp1 mRNA (Input)
Relative to Luciferase (2^-ΔCT) (Normalized to CM Ctrl.)
ns
ns
ns
ns
BCAAs + - + -
LPS - - + +
E
Casp1 mRNA (Fractions)
Relative to Luciferase (2^-ΔCT)
100
80
60
40
20
0
Complete Medium
BCAA-Free Medium
Complete Medium (+ LPS)
BCAA-Free Medium (+ LPS)
F4 F5 F6 F7 F8 F9 F10 F11 F12 F13
60S
Mono/Disomes
Polysomes
F
Casp1 mRNA Fraction Area
Polysome AUC (Relative to CM Ctrl.)
20
15
10
5
0
ns
p = 0.12
ns
ns
BCAAs + - + -
LPS - - + +

Figure S6

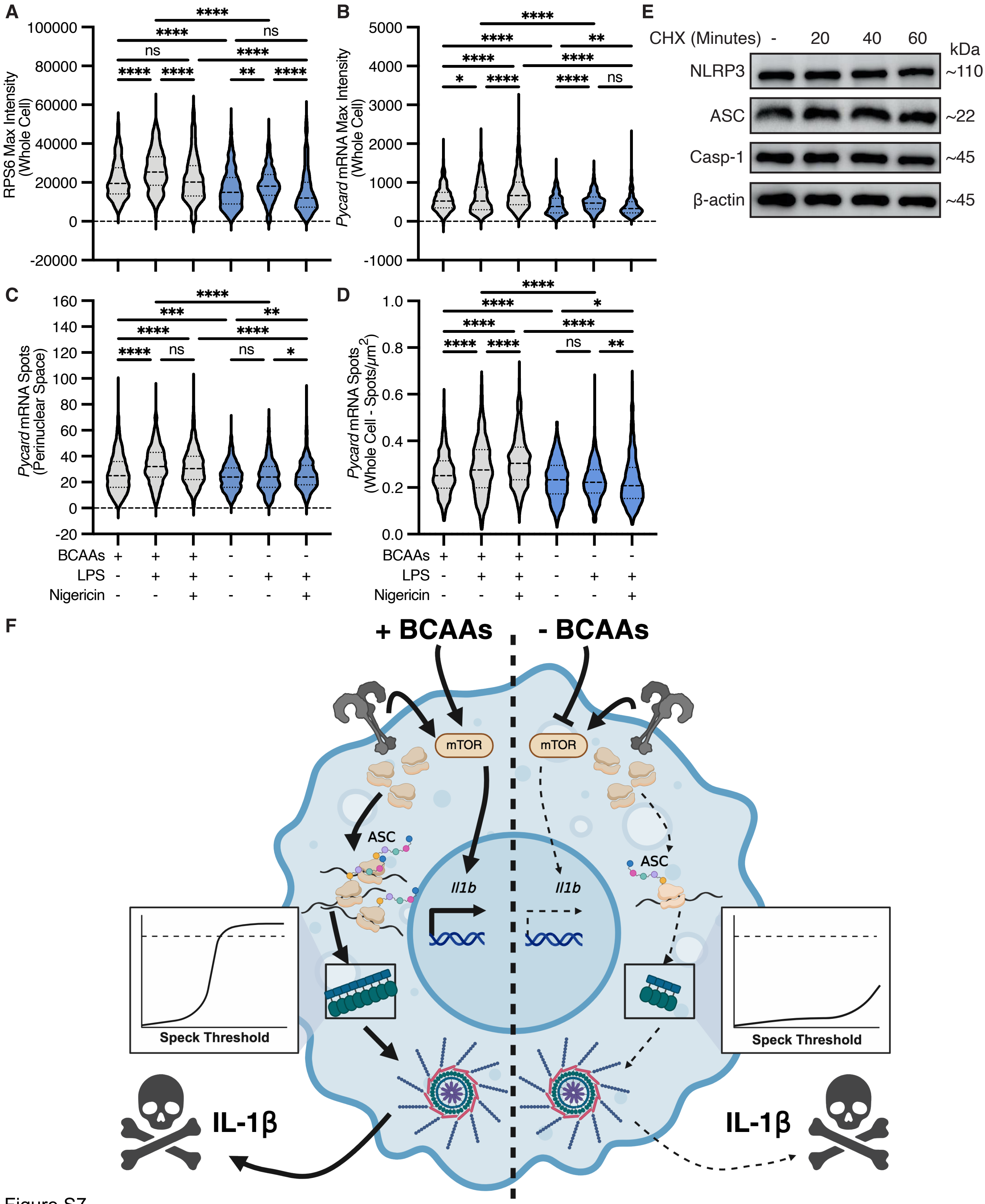

A
RPS6 Max Intensity (Whole Cell)
100000
80000
60000
40000
20000
0
-20000
****
****
ns
ns
****
****
****
**
****
B
Pycard mRNA Max Intensity (Whole Cell)
5000
4000
3000
2000
1000
0
-1000
****
****
**
****
****
*
****
****
ns
E
CHX (Minutes)
-
20
40
60
kDa
NLRP3
~110
ASC
~22
Casp-1
~45
β-actin
~45
C
Pycard mRNA Spots (Perinuclear Space)
160
140
120
100
80
60
40
20
0
-20
****
***
**
****
****
****
ns
ns
*
D
Pycard mRNA Spots (Whole Cell - Spots/μm2)
1.0
0.8
0.6
0.4
0.2
0.0
****
****
*
****
****
****
****
ns
**
BCAAs + + + - - -
LPS - + + - + +
Nigericin - - + - - +
F
+ BCAAs
- BCAAs
mTOR
mTOR
ASC
ASC
Il1b
Il1b
Speck Threshold
Speck Threshold
IL-1β
IL-1β


Figure S7

**AMINO ACID DROPOUT SCREEN**

**STEP #1**

**Preparation of Amino Acid-Free "Base" Medium**

RPMI 1640 Medium Modified w/o L-Glutamine, w/o Amino acids, Glucose (Powder) (United States Biological, #R9010-01)

*Prepared according to manufacturer's instructions and supplemented with 2 g/L glucose to generate an amino acid-free "base" medium (pH'd to 7.2).*

**STEP #2**

**Preparation of Amino Acid Stock Solutions**

*30 mL of each 100x amino acid stock solution were prepared in the base medium specified above. Stock solutions were then sterile-filtered and aliquoted (~5 mL in each aliquot) for long-term storage at -20°C.*

| Amino Acid | RPMI 1640 Concentration (mg/L) | 100x Concentration (mg/L) | mg in 30 mL for 100x | NOTES | |
|---|---|---|---|---|---|
| Arginine (Sigma-Aldrich, #A5006) | 200 | 20000 | 600 | | [redacted] |
| Asparagine (Sigma-Aldrich, #A7094) | 56.82 | 5682 | 170.46 | | |
| Aspartic Acid (Sigma-Aldrich, #A9256) | 20 | 2000 | 60 | | |
| Cystine (Sigma-Aldrich, #C6727) | 65.2 | 6520 | 195.6 | *Dissolved in base medium containing 1M HCl (i.e., 2.5 mL 12M HCl in 27.5 mL base medium)* | |
| Glutamic Acid (Sigma-Aldrich, #G8415) | 20 | 2000 | 60 | | |
| Glutamine (Sigma-Aldrich, #G8540) | 300 | 30000 | 900 | | |
| Glycine (Fisher BioReagents, #BP381) | 10 | 1000 | 30 | | |
| Histidine (Sigma-Aldrich, #H6034) | 15 | 1500 | 45 | | |
| Hydroxyproline (Cayman Chemical, #26061) | 20 | 2000 | 60 | | |
| Isoleucine (Sigma-Aldrich, #I2752) | 50 | 5000 | 150 | | |
| Leucine (Sigma-Aldrich, #L8000) | 50 | 5000 | 150 | | |
| Lysine (Sigma-Aldrich, #L5626) | 40 | 4000 | 120 | | |
| Methionine (Sigma-Aldrich, #M9625) | 15 | 1500 | 45 | | |
| Phenylalanine (Sigma-Aldrich, #P5482) | 15 | 1500 | 45 | | |
| Proline (Sigma-Aldrich, #P5607) | 20 | 2000 | 60 | | |
| Serine (Sigma-Aldrich, #S4311) | 30 | 3000 | 90 | | |
| Threonine (Sigma-Aldrich, #T8625) | 20 | 2000 | 60 | | |
| Tryptophan (Sigma-Aldrich, #T0254) | 5 | 500 | 15 | | |
| Tyrosine (Sigma-Aldrich, #T1145) | 28.83 | 2883 | 86.49 | | |
| Valine (Sigma-Aldrich, #V0500) | 20 | 2000 | 60 | | |

**STEP #3**

**Preparation of Amino Acid Dropout Media**

*Base medium was selectively supplemented with 100x amino acid stock solutions to generate complete and amino acid dropout media, as appropriate (see table embedded in "Pipetting Grid"). NaOH was then added to achieve physiologic pH before addition of dialyzed FBS.*

| Culture Medium | Total Volume (mL) | Volume Base Medium (mL) | Volume Stock Solutions | Volume Dialyzed FBS (10% Final) | Volume 5M NaOH | NOTES |
|---|---|---|---|---|---|---|
| Complete (i.e., all amino acids) | 30 mL | 21.6 mL | 20 x 270 uL = 5.4 mL | 3 mL | 1-10 uL | |
| - Arginine | 20 mL | 14.58 mL | 19 x 180 uL = 3.42 mL | 2 mL | 1-10 uL | |
| - Asparagine | 20 mL | 14.58 mL | 19 x 180 uL = 3.42 mL | 2 mL | 1-10 uL | |
| - Aspartatic Acid | 20 mL | 14.58 mL | 19 x 180 uL = 3.42 mL | 2 mL | 1-10 uL | |
| - Cystine | 20 mL | 14.4 mL + 0.18 mL base medium containing 1M HCl | 19 x 180 uL = 3.42 mL | 2 mL | 1-10 uL | *In place of 100x cystine, 0.18 mL base medium containing 1M HCl was added* |
| - Glutamatic Acid | 20 mL | 14.58 mL | 19 x 180 uL = 3.42 mL | 2 mL | 1-10 uL | |
| - Glutamine | 20 mL | 14.58 mL | 19 x 180 uL = 3.42 mL | 2 mL | 1-10 uL | |
| - Glycine | 20 mL | 14.58 mL | 19 x 180 uL = 3.42 mL | 2 mL | 1-10 uL | |
| - Histidine | 20 mL | 14.58 mL | 19 x 180 uL = 3.42 mL | 2 mL | 1-10 uL | |
| - Hydroxyproline | 20 mL | 14.58 mL | 19 x 180 uL = 3.42 mL | 2 mL | 1-10 uL | |
| - Isoleucine | 20 mL | 14.58 mL | 19 x 180 uL = 3.42 mL | 2 mL | 1-10 uL | |
| - Leucine | 20 mL | 14.58 mL | 19 x 180 uL = 3.42 mL | 2 mL | 1-10 uL | |
| - Lysine | 20 mL | 14.58 mL | 19 x 180 uL = 3.42 mL | 2 mL | 1-10 uL | |
| - Methionine | 20 mL | 14.58 mL | 19 x 180 uL = 3.42 mL | 2 mL | 1-10 uL | |
| - Phenylalanine | 20 mL | 14.58 mL | 19 x 180 uL = 3.42 mL | 2 mL | 1-10 uL | |
| - Proline | 20 mL | 14.58 mL | 19 x 180 uL = 3.42 mL | 2 mL | 1-10 uL | |
| - Serine | 20 mL | 14.58 mL | 19 x 180 uL = 3.42 mL | 2 mL | 1-10 uL | |
| - Threonine | 20 mL | 14.58 mL | 19 x 180 uL = 3.42 mL | 2 mL | 1-10 uL | |
| - Tryptophan | 20 mL | 14.58 mL | 19 x 180 uL = 3.42 mL | 2 mL | 1-10 uL | |
| - Tyrosine | 20 mL | 14.58 mL | 19 x 180 uL = 3.42 mL | 2 mL | 1-10 uL | |
| - Valine | 20 mL | 14.58 mL | 19 x 180 uL = 3.42 mL | 2 mL | 1-10 uL | |

Table S1

| Culture Medium | Arg | Asn | Asp | Cys | Glu | Gln | Gly | His | H-Pro | Ile | Leu | Lys | Met | Phe | Pro | Ser | Thr | Trp | Tyr | Val |
|---|---|---|---|---|---|---|---|---|---|---|---|---|---|---|---|---|---|---|---|---|
| **Complete (i.e., all amino acids)** | x | x | x | x | x | x | x | x | x | x | x | x | x | x | x | x | x | x | x | x |
| **- Arginine** | | x | x | x | x | x | x | x | x | x | x | x | x | x | x | x | x | x | x | x |
| **- Asparagine** | x | | x | x | x | x | x | x | x | x | x | x | x | x | x | x | x | x | x | x |
| **- Aspartatic Acid** | x | x | | x | x | x | x | x | x | x | x | x | x | x | x | x | x | x | x | x |
| **- Cystine** | x | x | x | | x | x | x | x | x | x | x | x | x | x | x | x | x | x | x | x |
| **- Glutamatic Acid** | x | x | x | x | | x | x | x | x | x | x | x | x | x | x | x | x | x | x | x |
| **- Glutamine** | x | x | x | x | x | | x | x | x | x | x | x | x | x | x | x | x | x | x | x |
| **- Glycine** | x | x | x | x | x | x | | x | x | x | x | x | x | x | x | x | x | x | x | x |
| **- Histidine** | x | x | x | x | x | x | x | | x | x | x | x | x | x | x | x | x | x | x | x |
| **- Hydroxyproline** | x | x | x | x | x | x | x | x | | x | x | x | x | x | x | x | x | x | x | x |
| **- Isoleucine** | x | x | x | x | x | x | x | x | x | | x | x | x | x | x | x | x | x | x | x |
| **- Leucine** | x | x | x | x | x | x | x | x | x | x | | x | x | x | x | x | x | x | x | x |
| **- Lysine** | x | x | x | x | x | x | x | x | x | x | x | | x | x | x | x | x | x | x | x |
| **- Methionine** | x | x | x | x | x | x | x | x | x | x | x | x | | x | x | x | x | x | x | x |
| **- Phenylalanine** | x | x | x | x | x | x | x | x | x | x | x | x | x | | x | x | x | x | x | x |
| **- Proline** | x | x | x | x | x | x | x | x | x | x | x | x | x | x | | x | x | x | x | x |
| **- Serine** | x | x | x | x | x | x | x | x | x | x | x | x | x | x | x | | x | x | x | x |
| **- Threonine** | x | x | x | x | x | x | x | x | x | x | x | x | x | x | x | x | | x | x | x |
| **- Tryptophan** | x | x | x | x | x | x | x | x | x | x | x | x | x | x | x | x | x | | x | x |
| **- Tyrosine** | x | x | x | x | x | x | x | x | x | x | x | x | x | x | x | x | x | x | | x |
| **- Valine** | x | x | x | x | x | x | x | x | x | x | x | x | x | x | x | x | x | x | x | |

*x = added to medium*

Table S1 - Pipetting Grid
